# Oxphos Targeting of *Mycn*-Amplified Neuroblastoma

**DOI:** 10.1101/2024.08.03.606365

**Authors:** Soraya Epp, Donagh Egan, Evon Poon, Amirah Adlina Abdul Aziz, Kieran Wynne, Louis Chesler, Melinda Halasz, Walter Kolch

## Abstract

**Summary:** High risk - neuroblastoma (HR-NB) is a pediatric solid tumor with high lethality. Half of HR-NB are driven by *MYCN* gene amplification (MNA). These HR-NBs require high dosage chemotherapy and often relapse. Moreover, current therapies can cause severe long-term side effects and new therapies are urgently needed. This study investigates a novel therapeutic approach targeting the metabolic vulnerabilities of MNA NB cells. We discovered that Diphenyleneiodonium chloride (DPI), an inhibitor of flavoprotein enzymes and mitochondrial complex I, synergizes with mitoquinone mesylate (MitoQ), a mitochondria-targeted antioxidant in 2D and 3D *in vitro* models of NB. Similarly to DPI, MitoQ affects MNA cells in a MYCN-dependent fashion, being more toxic when MYCN levels are high. Furthermore, low nanomolar concentrations of MitoQ significantly decrease MYCN protein expression and induce differentiation of MNA cells. The DPI and MitoQ combination further synergizes with vincristine, a chemotherapeutic agent used in NB treatment. Phosphoproteomics and proteomics analysis suggests that the drug combination induces MNA NB cell death by arresting the cell cycle and inhibiting oxidative phosphorylation (OXPHOS) in the mitochondria. Thus, interference with mitochondrial metabolism may represent an effective strategy to enhance the activity of chemotherapeutic drugs in MNA-NB.

## 1. Introduction

Neuroblastoma (NB) is the most common extracranial solid tumor in infants. NBs originate from neural crest cells and typically form in the abdominal or thoracic regions in paraspinal ganglia^1^. Despite comprising only 6% of all pediatric cancers, NB accounts for 15% of all childhood cancer-related deaths^2^. NB is unique due to its clinical bipolarity: favorable with occasional spontaneous regression, or unfavorable, with highly aggressive behavior and metastatic spread^1,3^. Despite considerable progress in improving survival rates, the overall 5-year event-free survival for high-risk patients (ca. 40% of all NB patients) is still less than 50%. High-risk patients are often treated with a combination of intensive genotoxic chemotherapy and radiation therapy. However, a large proportion of patients relapses, leaving them with limited or no treatment options. Furthermore, genotoxic therapies can result in significant long-term side effects, such as hearing loss, secondary cancers, infertility, and endocrine deficiencies^4^.

MYCN is a transcription factor belonging to the highly homologous *MYC*-family of cellular oncogenes, ubiquitously expressed in mammals. *MYCN* amplification (MNA) occurs in ca. 20% of NB patients and drives half of the aggressive forms of the disease with poor outcomes^1^. Pharmacologic inhibition of MYCN is very challenging due to the absence of drug-binding pockets in the MYCN protein, and no FDA-approved treatment is currently available^5^. Thus, efforts have focused on strategies to block MYCN indirectly, e.g. via inhibition of the *MYCN* transcriptional machinery, synthetic lethality, and destabilization of the MYCN protein ^6,7^.

NB, like other malignancies, is associated with modifications in cellular metabolism that may contribute to oncogenesis. Activation of oncogenes and loss of tumor suppressors lead to the increased production of superoxide via mitochondria and NADPH oxidase (NOX) pathways^8,9^. Superoxide is subsequently converted to hydrogen peroxide (H_2_O_2_), a more stable and stronger oxidizing molecule, through the action of superoxide dismutase enzymes (SOD). H_2_O_2_ can oxidize proteins and lipids, but also act as a signaling molecule modulating various cellular processes linked to oncogenesis^10^. As a result, malignant cells need to control the concentration of H _2_O_2_ precisely to avoid excessive oxidative damage, while enabling the pro-tumorigenic effects of H_2_O_2_ signaling. MNA cells rely on an elevated antioxidant response through the Xc-/glutathione (GSH) pathway to scavenge lipid hydroperoxides, a by-product formed from excess H_2_O_2_^11,12^. The preference for using oxidative phosphorylation (OXPHOS) to meet energy demands, and the resulting increase in reactive oxygen species (ROS) has been shown to be a targetable vulnerability of MNA cells^8,12,13^. Recently, several studies have demonstrated the ability of MYCN to regulate mitochondrial genes in NB, and more specifically complex I genes of the electron transport chain (ETC)^14,15^.

We recently discovered in a drug screen that Diphenyleneiodonium chloride (DPI) selectively induces cell death of MNA NB cells *in vitro* and decreases tumor growth *in vivo*^16^. DPI is an inhibitor of flavoprotein enzymes, including NOX, and the mitochondrial complex I. It has a wide spectrum of metabolic effects, including induction of apoptosis via ROS production and cytochrome c release^17^, inhibition of NADH oxidation^18^, and loss of GSH^19^. DPI has also been employed in numerous studies as a ROS inhibitor due to its demonstrated efficacy in reducing superoxide production during reverse electron transport^20,21^. However, identifying the specific ROS generation site and the precise mode of DPI action has proven challenging and appears to be contingent upon contextual factors^22,23^. We also demonstrated that DPI significantly increases mitochondrial ROS production in MNA cells, but not in non-MNA cells, indicating a MYCN-dependent effect^16^. To investigate the role of ROS and the mechanism of action of DPI in MNA cells, we employed a widely and commercially available mitochondria-targeted antioxidant, mitoquinone mesylate (MitoQ). Interestingly, we found that DPI synergizes with MitoQ. Therefore, we evaluated the combined effects of DPI, MitoQ and vincristine, a chemotherapeutic drug in use for the treatment of high-risk NB.

Although precision therapy for individual patients remains limited, combination therapy has the potential to enhance treatment efficacy while potentially reducing reliance on highly toxic chemotherapy^24^. Multiple drug combinations have several advantages over monotherapy, as they can achieve higher efficacy and, by using lower drug doses, reduce the risk of adverse effects and acquired resistance^25^. As a result, combination therapies are widely utilized in the treatment of cancer^26^, although primarily involving genotoxic drugs for the treatment of NB. To address this issue, we evaluated a potential triple combination therapy, comprising two metabolic-targeting drug and a chemotherapeutic agent, aiming to develop a more effective therapeutic approach for MNA NB.

## 2. Results

### DPI in combination with MitoQ induces cell death and differentiation of MNA NB cells

To investigate whether enhanced ROS production mediates the cytotoxic effects of DPI, we used MitoQ, a ROS scavenger that selectively accumulates in mitochondria^27,28^. First, we assessed the viability of the MNA IMR-5/75 cells in response to MitoQ. In these cells MYCN protein expression can be downregulated by ca. 50% by a doxycycline inducible shRNA^29^ (**Fig. S1**). We found that MitoQ was non-toxic up to a concentration of 100 nM over a 24-hour treatment period (**Fig. 1A**). Then, we tested the antioxidant effect of MitoQ when combined with DPI. Mitochondrial superoxide production was measured using the selective fluorogenic MitoSOX Red dye^30^. DPI is known to increase mitochondrial superoxide production in a dose dependent manner^16^. Unexpectedly, MitoQ on its own increased ROS concentrations, although the effect was not significant (**Fig. 1B**). Combining DPI and MitoQ effectively suppressed DPI-induced superoxide production to almost baseline level (**Fig. 1B**). To determine whether cell death follows the pattern of superoxide levels recorded in DPI and MitoQ-treated IMR-5/75 cells, cell toxicity was measured using the CellTox Green Dye that detects modifications in membrane integrity resulting from cell death (**Fig. 1A, C**). As previously shown, DPI induced progressive cell death over the three-day observation period, which was significantly mitigated by downregulation of MYCN^31^ (**Fig. 1C**). 100 nM MitoQ on its own was significantly cytotoxic at day 2 and 3, an effect which was also lessened by downregulation of MYCN (**Fig. 1A, C**). However, MitoQ failed to reduce DPI-induced cell death at all tested time points (**Fig. 1C**). Cell death was further assessed by counting nuclei and Propidium Iodide (PI) staining (**Fig. 1D, 1E**). Nuclei counting of IMR-5/75 cells treated with the same conditions (**Fig. 1D**) confirmed the cell toxicity results (**Fig. 1C**). DPI strongly reduced the number of nuclei indicating loss of viable cells. MitoQ addition to DPI significantly reduced the numbers of nuclei after 72 hours. PI cannot penetrate intact cell membranes and thus only stains cells whose membrane integrity is compromised, such as apoptotic or necrotic cells. Interestingly, neither high dose DPI nor 1 μM MitoQ induced PI positive cells significantly on their own after 24 hours of treatment. While 100 nM MitoQ did not affect DPI-induced cell death (**Fig. 1C**), higher MitoQ concentrations significantly enhanced DPI-mediated cell toxicity as measured by an increased PI uptake, in a MYCN-dependent fashion (**Fig. 1E**).

**Fig 1.**
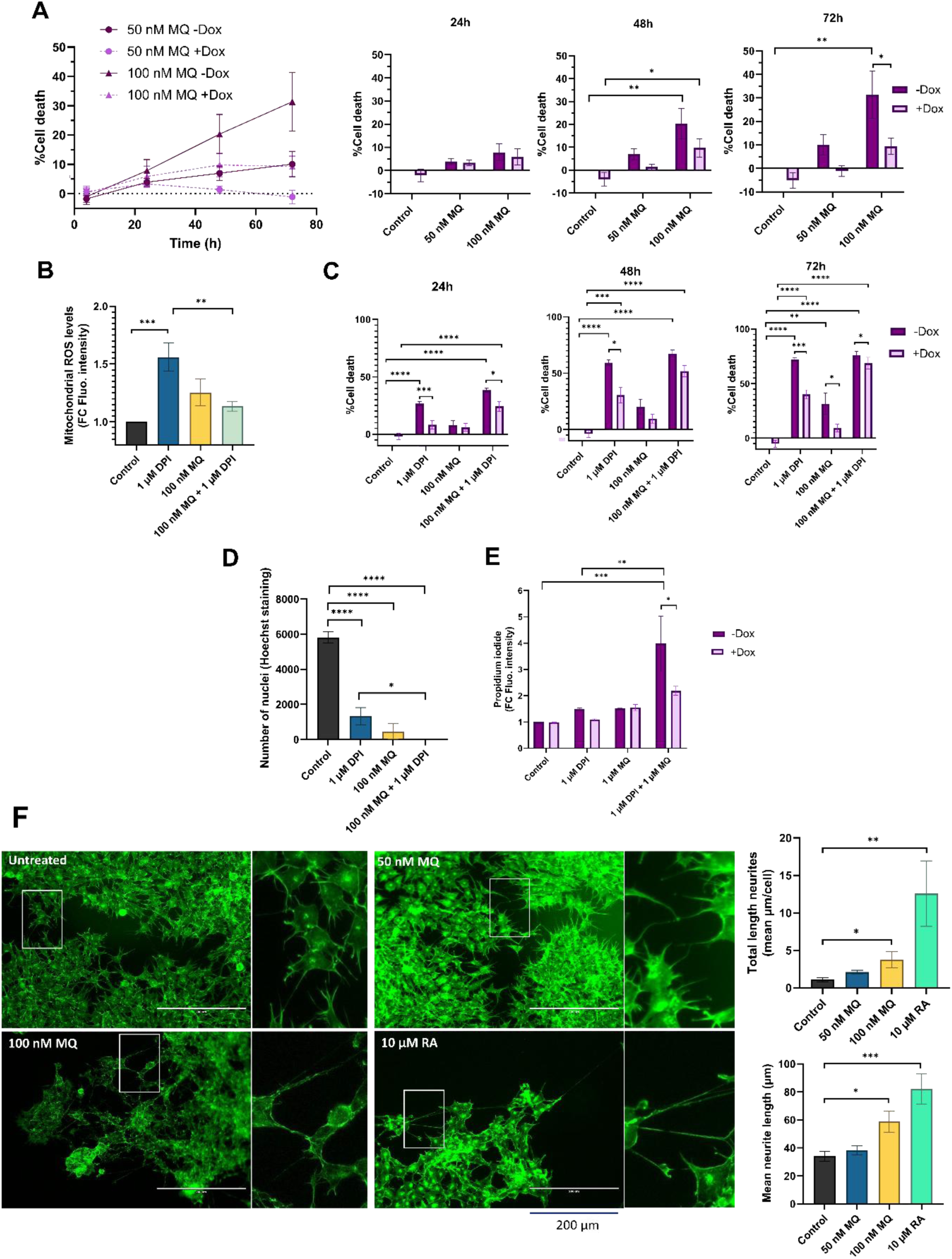
DPI in combination with MitoQ induces cell death and differentiation of NB cells. **A)** CellTox Green dye was used to determine cytotoxicity of IMR-5/75 cells -/+ Dox treated with MitoQ over a period of 72 hours. +Dox: shRNA targeting MYCN induced and MYCN expression reduced; -Dox: shRNA targeting MYCN not induced. (Mean ± SEM; n=3; Two-Way ANOVA, p<0.05 = *, p<0.005 = **). **B)** MitoSOX Red dye was used to measure mitochondrial superoxide production in IMR-5/75 treated with DPI and MitoQ alone or in combination for 2 hours. (Mean ± SEM; n=3; Ordinary One-Way ANOVA, p<0.05 = *, p<0.005 = **, p<0.0005 = ***). **C)** CellTox Green dye was used to determine cytotoxicity of IMR-5/75 cells -/+ Dox treated with DPI and MitoQ alone or in combination over a period of 72 hours. (Mean ± SEM; n=3; Two-Way ANOVA, p<0.05 = *, p<0.005 = **, p<0.0005 = ***, p<0.0001 = ****). **D)** Nuclei of IMR-5/75 cells were visualized by Hoechst staining after 72 hours of treatment with either DPI or MitoQ on their own or in combination. (Mean ± SEM; n=3; Ordinary One-Way ANOVA, p<0.05 = *, p<0.0001 = ****). **E)** Propidium Iodide staining was observed by confocal microscopy to quantify cell death in IMR-5/75 following a 24-hour treatment with DPI, MitoQ or a combination of both in -Dox and +Dox cells. (Mean ± SEM; n=3; Two-Way ANOVA, p<0.05 = *, p<0.005 = **, p<0.0005 = ***, p<0.0001 = ****). **F)** Neuronal differentiation was assessed by phalloidin staining of IMR-5/75 cells treated with 50 nM or 100 nM MitoQ for 7 days. 10 μM retinoic acid was used as a positive control to induce neurite outgrowth. Were taken into consideration cells with neurites >= 2 cell body diameters. Scale bar = 200 μm. (Mean ± SEM; n=3; Ordinary One-Way ANOVA, p<0.05 = *, p<0.005 = **, p<0.0005 = ***).

These results suggest that the cytotoxicity observed following treatment with MitoQ alone and in combination with DPI is not solely dependent on its ability to regulate mitochondrial superoxide production in MNA NB cells, which is in line with recent studies on MitoQ effects in other cancer cell types^32,33^. Interestingly, the cytotoxic effects of MitoQ were amplified by higher MYCN levels.

The arrested differentiation of neuroblasts is a characteristic of NB tumorigenesis and invasiveness^34^. To assess whether MitoQ treatment can similarly overcome the differentiation block observed with DPI^16^, MNA IMR-5/75 cells were treated every two days with either 50 nM or 100 nM MitoQ and monitored by microscopy (**Fig. 1F**). Neurite outgrowth was visualized and quantified using Phalloidin staining, which specifically binds to actin filaments. 100 nM MitoQ treatment resulted in differentiation as evidenced by an increase in neurite outgrowth 7 days after treatment, compared to the non-treated control. 10 μM retinoic acid was used as a positive control for maturation towards a differentiated neuronal phenotype, as shown previously^35^. These results suggest that MitoQ slows down cell growth, induces cell death and differentiation of MNA NB cells.

### DPI synergizes with MitoQ in MNA NB cells

To assess whether MitoQ and DPI act synergistically in reducing cell viability, we treated two MNA NB cell lines (IMR-5/75 and Be(2)-C) with various concentrations of DPI and MitoQ. Synergy was assessed using SynergyFinder 3.0^36^, which computes synergy scores based on different mathematical models. Given that both MitoQ and DPI target mitochondrial function, we used the Loewe synergy model, which is particularly suited for evaluating the combined effects of drugs with similar mechanisms of action^37^. The combination exhibited a pronounced synergistic effect in both MNA NB cell lines (**Fig. 2A, B**). Although the concept of combination synergy is highly dependent on the underlying assumptions^38^, according to SynergyFinder 3.0 the synergy scores can be interpreted as follows : from -10 to 10 the interaction is likely to be additive, larger than 10 the interaction is likely to be synergistic, and conversely, if the score in less than -10, then the interaction between the two drugs is most likely antagonistic. In IMR-5/75, the highest Loewe score is 43.2 and obtained at ∼400 nM MitoQ combined with ∼500 nM DPI (**Fig. 2A**). The highest Loewe score in Be(2)-C is 45.3 and reached at ∼1800 nM MitoQ combined with ∼10 nM DPI (**Fig. 2B**). The highest score was reached with a ∼10fold lower concentration of MitoQ in IMR-5/75 than in Be(2)-C cells, which seem to be more resistant to treatment (**Fig. 2A, B**). These results are in line with previous findings indicating that inhibition of mitochondrial complex I synergizes with mitochondrial antioxidants in impairing cellular fitness of cancer cells^28^.

**Fig 2.**
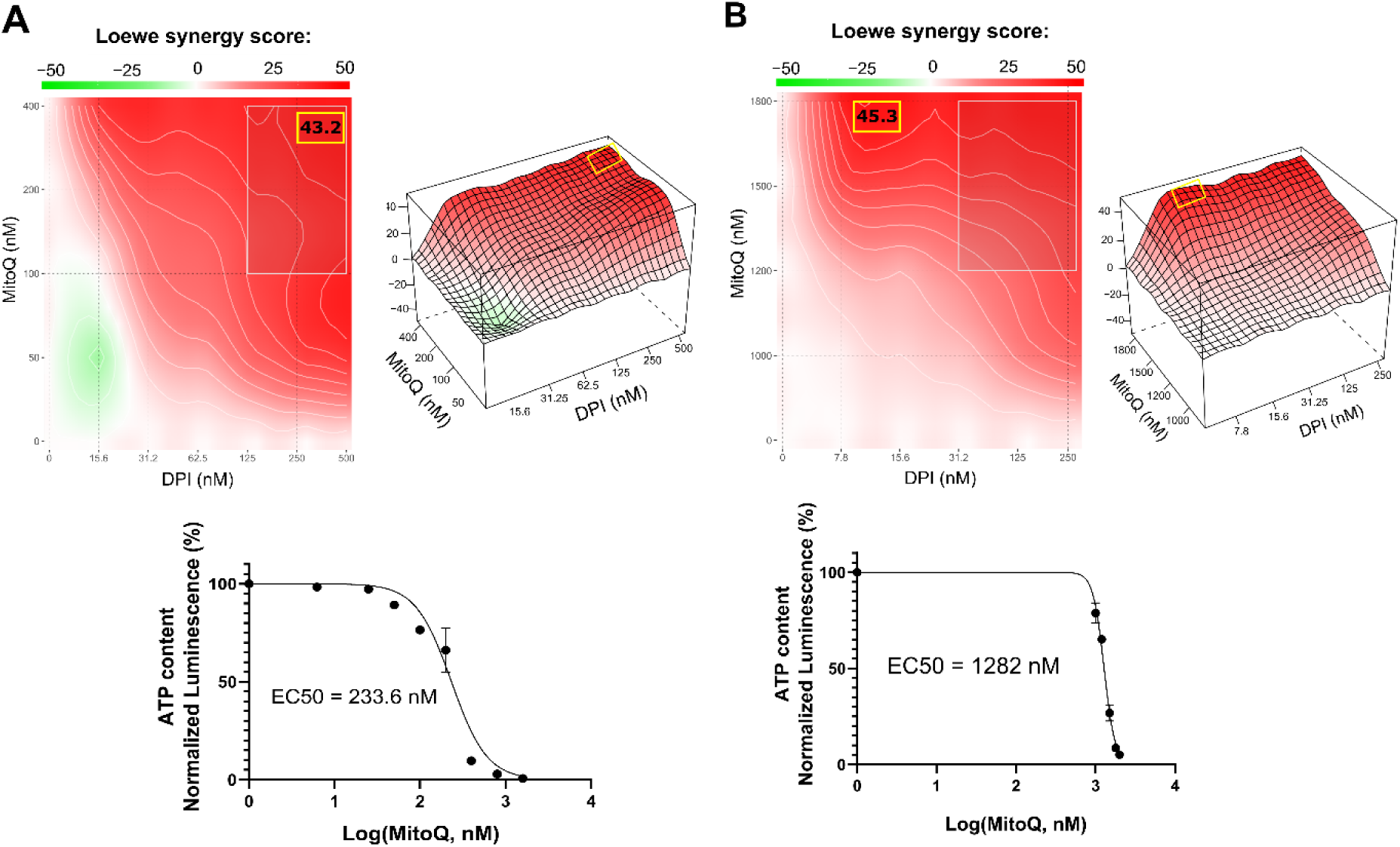
DPI synergizes with MitoQ in MNA NB cells. SynergyFinder 3.0 analysis was used to generate Loewe synergy heatmaps, generated from cell viability values (CellTiter-Glo) of the combination of DPI with MitoQ, in **A)** IMR-5/75, and **B)** Be(2)-C. (Mean ± SEM; n=3). Lower panels: corresponding drug-response curves were generated from cell viability values after 72 hours of treatment. (Mean ± SEM; n>3).

### DPI synergizes with vincristine and MitoQ in reducing cell viability of MNA NB cells

Finding synergistic drug combinations allows for combining drugs at lower concentrations, which is an important strategy for enhancing therapeutic efficacy while minimizing adverse effects. Vincristine (vin) is a commonly used chemotherapeutic agent for treating patients with NB. As genotoxic treatment is clinically very effective but also toxic, we tested whether DPI could reduce the dosage of vincristine required for efficient cancer cell killing. A DPI and vincristine combination was tested for synergy in the two MNA NB cell lines IMR-5/75 and Be(2)-C (**Fig. 3A, B**). To evaluate this combination, we used the Zero Interaction Potency (ZIP) model, which accounts for independent drug effects and is particularly suited for combinations of agents with differing targets, such as DPI and vincristine^39^. ZIP scores of >10 are considered to indicate synergy^36^. The combination showed synergistic effects in reducing cell viability in IMR-5/75 and Be(2)-C cells. In IMR-5/75, the highest ZIP score is 11.3 and reached at ∼0.75 nM vincristine combined with ∼100 nM DPI (**Fig. 3A**). The highest ZIP score in Be(2)-C is 13.7 and observed for ∼15 nM vincristine combined with ∼30 nM DPI (**Fig. 3B**). These results indicate that vincristine and DPI have an additive/synergistic interaction in MNA cells for reducing cell viability at low concentrations that are clinically achievable.

**Fig 3.**
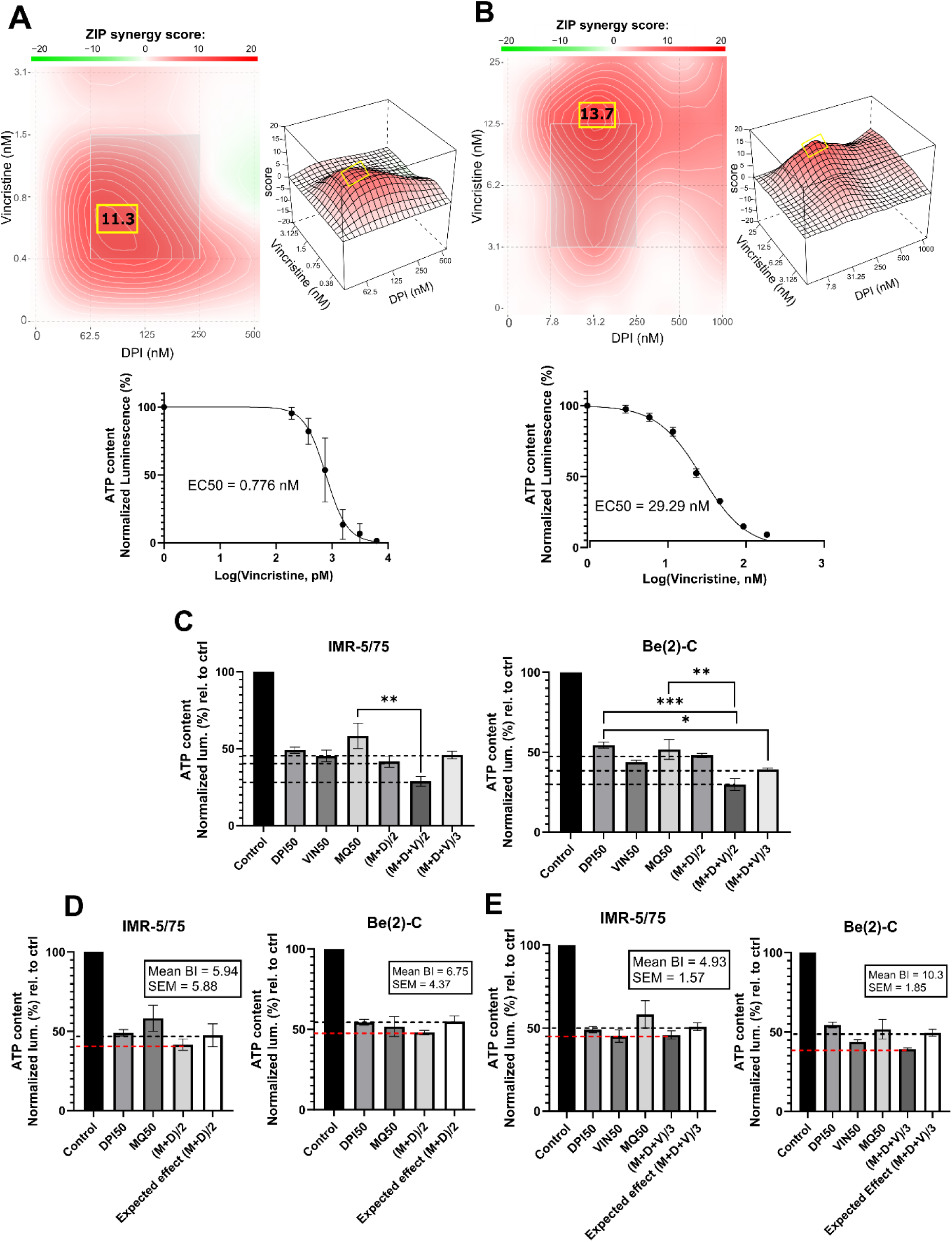
DPI synergizes with vincristine and MitoQ in reducing cell viability of MNA NB cells. SynergyFinder 3.0 was used to generate ZIP synergy heatmaps from cell viability values (CellTiter-Glo) of the combination of DPI with vincristine, in **A)** IMR-5/75, and **B)** Be(2)-C after 72 hours of treatment. Lower panels, corresponding drug-response curves generated from vincristine-treated cell viability values to determine their EC50 concentrations. (Mean ± SEM; n>3). **C)** IMR-5/75 and Be(2)-C cells were treated with reduced EC50 doses of MitoQ, DPI and vincristine. (M = MitoQ, D = DPI, and V = vincristine) (Mean ± SEM; n=3; Ordinary One-Way ANOVA, p<0.05 = *, p<0.005 = **, p<0.0005 = ***). **D, E)** The Bliss Independence model was used to assess synergy, additivity or antagonism of the double and triple drug combination at a reduced dose in IMR-5/75 and Be(2)-C MNA cells. Dashed black lines indicate the expected effect, and the dashed red lines the observed effect. (Mean ± SEM; n=3).

Given the synergy of DPI with both MitoQ and vincristine, we combined all three drugs at reduced dosage. To assess synergy of the 3-drug combination, the Bliss Independence model was used^40^ (**Fig. 3C, D, E**). The model is based on the premise that the relative effect of a drug is independent of the presence of the other drug. When two or more independent drugs are combined, their effects add up, and the Bliss synergy score is 0^40^. Within the Bliss independence model calculation, synergy is assumed when the actual effect of the combination is greater than the expected effect, i.e., if the Bliss score is greater than 0^41^. First, the EC50 of each drug was determined in two MNA cell lines (Be(2)-C and IMR-5/75) using the CellTiter-Glo assay (Promega) (**Fig. 3A, lower panels**). Then, for a 2-drug combination (MitoQ + DPI, (M+D)/2), the drugs were tested at their EC50 concentration divided by 2 or for a 3-drug combination (MitoQ + DPI + vin, (M+D+V)/3) divided by 3. For both MNA NB cell lines tested, the EC50/2 and EC50/3 combinations displayed similar or slightly greater effects in reducing cell viability than the EC50 concentration of each of the single drugs. Indeed, for all the combinations, the Bliss score was > 0 in the two MNA NB cell lines **(Fig. 3D, E)**. Therefore, the double and triple combinations at reduced doses are synergistic in reducing the cell viability of IMR-5/75 and Be(2)-C.

### Combining DPI, MitoQ and vincristine induces cell death of MNA NB cells

To characterize the interaction between MitoQ, DPI, and vincristine, we assessed their effects on cell death at EC50/2 and EC50/3 concentrations in four NB cell lines: Be(2)-C (MNA), IMR-32 (MNA), IMR-5/75 (MNA with MYCN downregulatable), and SH-SY5Y/MYCN (non-MNA, MYCN-inducible). The Be(2)-C cell line showed greater resistance with EC50s higher than other tested cell lines. Therefore, in order to assure that the drug sensitivity range is covered in full, the EC50 concentrations of Be(2)-C cells for the 3 drugs were taken as a basis to treat all other cell lines. Cell toxicity was measured using CellTox Green (**Fig. 4**). The double combination ((M+D)/2) significantly increased toxicity in IMR-32 and IMR-5/75 cells within 24 hours, persisting up to 72 hours. However, in Be(2)-C cells, it reduced ATP content by more than half (**Fig. 3D**) without increasing cell death (**Fig. 4A**), indicating a cytostatic rather than cytotoxic effect. In Be(2)-C, significant toxicity occurred only after 72 hours with the triple combination ((M+D+V)/3) (**Fig. 4C**). In SH-SY5Y/MYCN cells, (M+D)/2 had no effect on cell death, while (M+D+V)/3 increased toxicity at 48 and 72 hours (**Fig. 4A**). MYCN abundance modestly influenced sensitivity: MYCN downregulation in IMR-5/75 cells slightly reduced toxicity, while MYCN induction in SH-SY5Y/MYCN slightly increased it, though neither change reached significance. Although MYCN downregulation or overexpression are in the region of 50% or more, they are well within the natural variation of MYCN expression observed in MNA and MYCN non-amplified cells, respectively^42,43^. Overall, the triple-drug combination was effective in inducing cell death across four high-risk NB cell lines. The double-drug combination promoted cell death in two MNA cell lines, but in the more resistant MNA Be(2)-C cells it primarily caused metabolic inactivity.

**Fig 4.**
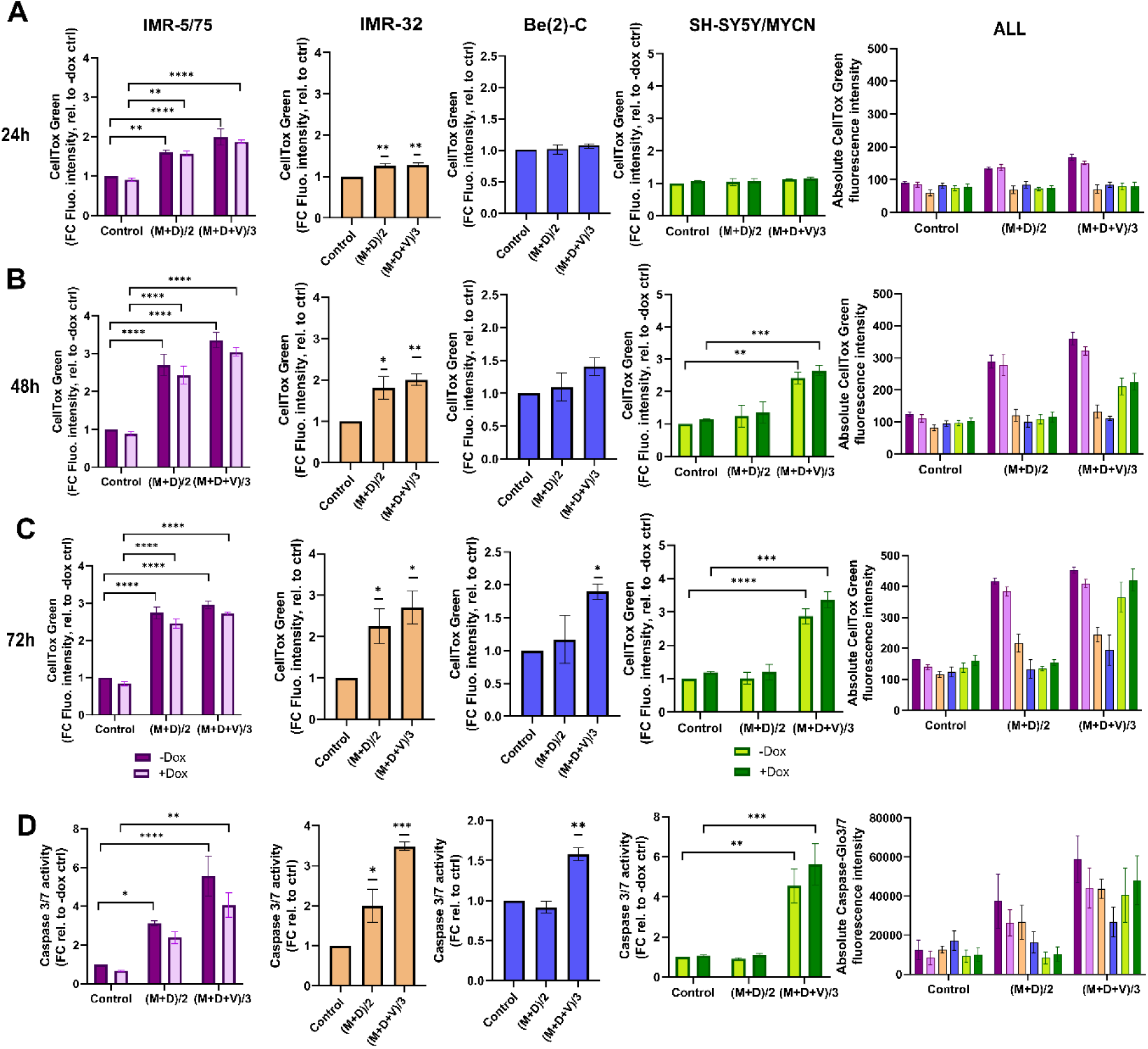
Double and triple combination therapy sensitizes HR-NB cells to cell death. **A-C)** CellTox Green dye or **D)** Caspase-Glo 3/7 was used to determine cytotoxicity and caspases 3/7 activation in IMR-5/75 cells -/+ Dox, IMR-32, Be(2)-C and SH-SY5Y/MYCN -/+ Dox with the double (DPI and MitoQ) or the triple (DPI and MitoQ and vincristine) combination after **A)** 24h, **B)** 48h and **C)** 72h of treatment. Caspases activation was measured at 24h. (Mean ± SEM; n=3; Ordinary One-Way ANOVA or Two-Way ANOVA for -/+ Dox cell lines, p<0.05 = *, p<0.005 = **, p<0.0005 = ***, p<0.0001 = ****). In the non-MYCN-amplified SH-SY5Y/MYCN cells, the expression of MYCN is doxycycline-inducible (+Dox: expression of the MYCN transgene induced; -Dox: MYCN transgene not induced).

To further measure cell death upon treatment with MitoQ, DPI and vincristine, we measured caspase 3/7 activation using Caspase-Glo 3/7 (Promega) (**Fig. 4D**). Caspases 3 and 7, key mediators of the mitochondrial apoptotic pathway, were significantly activated by (M+D)/2 after 24 hours in IMR-5/75 -Dox and IMR-32, but not in Be(2)-C and SH-SY5Y/MYCN cells. (M+D+V)/3 enhanced caspases 3/7 activation in all cell lines tested. Downregulating MYCN in IMR-5/75 decreased caspase activation for both treatments, while inducing MYCN expression in SH-SY5Y/MYCN increased caspase activation, although the differences were not statistically significant (**Fig. 4D**). These data suggest that MNA NB cells exhibit great sensitivity to the triple drug combinations. The non-MNA cell line SH-SY5Y/MYCN showed no sensitivity to the double combination, similarly to the most resistant MNA cell line, Be(2)-C. Combining compounds having a synergistic interaction enables the use of reduced concentrations while still inducing cell death and caspase activation at a high level.

### Combination therapy inhibits the growth of Th-*MYCN* mouse NB spheroids

To assess the antitumor potential of the dual and triple drug combinations in a more physiological model than cell lines, we used a 3D spheroid model of a transgenic mouse that overexpresses *MYCN* under the control of the tyrosine hydroxylase (TH) promoter. This promoter is active in neural crest cells, and driving *MYCN* overexpression in these progenitor cells of NB causes the animals (Th-*MYCN*) to develop tumors that closely resemble human NB^44^. Spheroid tumor models are 3D aggregates of cancer cells which mimic the architecture and cellular heterogeneity of tumor tissue *in vitro*. They are viewed as an intermediate model between 2D monolayers and animal models, as they preserve key features of tumors, especially the response to drug treatments^45^.

Treating the Th-*MYCN* spheroids with the DPI-MitoQ combination showed a strong synergistic effect. The highest Loewe score was 45.7, achieved when ≈ 300 nM MitoQ is combined with ≈ 7.8 nM DPI (**Fig. 5A**). The combination between DPI and vincristine was also synergistic but less potent. Peak synergy occurred when ≈ 2 nM vincristine was combined with 7.8 nM DPI, with an associated ZIP score of 13.6 (**Fig. 5B**). Interestingly, the synergy maps for both MitoQ-DPI and vincristine-DPI are highly similar to those observed in Be(2)-C cells. Moreover, the EC50 values of MitoQ, DPI and vincristine are within the range observed for IMR-5/75 and Be(2)-C MNA cell lines (**Fig. 2A,B**; **Fig. 3A, B**). These correlations show that 2D and 3D culture models of MNA behave very similarly in terms of drug responses, which validates drug screening in 2D cultures as not only as convenient but also meaningful method for finding new therapies for MNA.

**Fig. 5.**
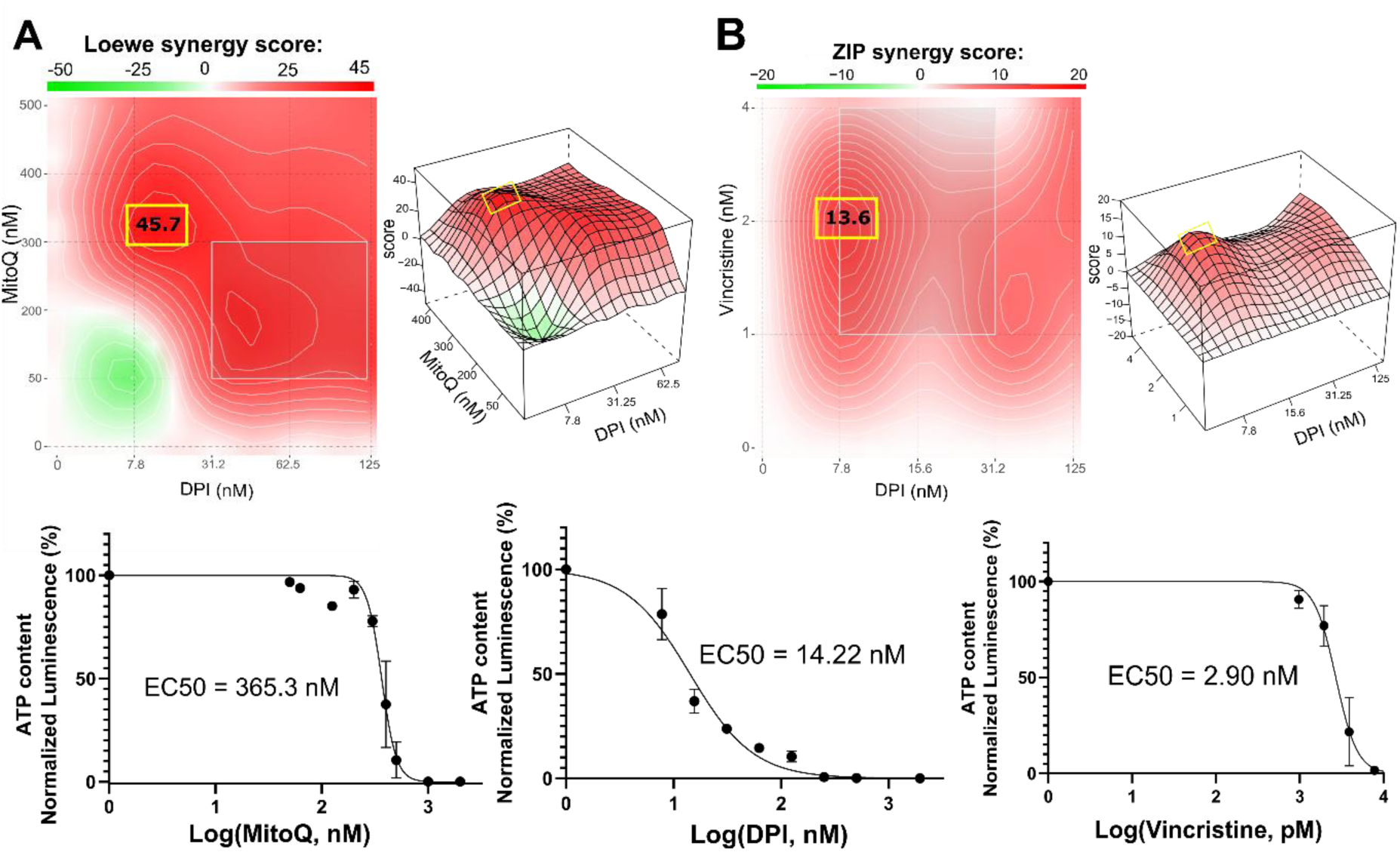
Combination therapy inhibits the growth of Th-MYCN mouse NB spheroids. SynergyFinder 3.0 analysis was used to generate Loewe and ZIP synergy heatmaps, generated from cell viability values (CellTiter-Glo) of the combination of **A)** DPI with MitoQ, and **B)** DPI with vincristine in Th-MYCN spheres. (Mean ± SEM; n=3). Below, associated drug-response curves generated from cell viability values after 72 hours of treatment. (Mean ± SEM; n>3).

In addition to assessing cell viability, DNA damage was visualized by PI^46,47^ and Hoechst staining of Th-*MYCN* spheres following exposure to M+D and M+D+V combinations (**Fig. S2**). The drugs were used at their EC50 concentrations to maximize synergy and induce a high rate of cell death. After 24 hours, both the double and the triple combinations increased cell death. The visual increase in Hoechst fluorescence intensity following M+D and M+D+V treatments can be explained by accumulation of the dye due to highly condensed and fragmented DNA ^47^. This observation confirms that the corresponding increase in PI intensity is due to apoptotic processes (**Fig. S2**).

### Investigating the mechanism of drug interaction by proteomic and phosphoproteomic analysis

To elucidate the synergistic mechanism of our double and triple drug combinations, we examined the proteomics and phosphoproteomics changes in MNA Be(2)-C cells treated with DPI+/-MitoQ+/-Vin for 24 hours by mass spectrometry (MS). The Be(2)-C cell line was selected due to its similar response profile as spheroids and delayed response to treatment compared to both MNA NB cell lines IMR-5/75 and IMR-32. Indeed, Be(2)-C cells only significantly apoptose after 72 hours of (M+D+V)/3 treatment (**Fig. 4C**), allowing a broad time window to identify pathways that mediate apoptosis-associated signaling.

First, we assessed protein expression in Be(2)-C cells treated with either MitoQ EC50, DPI EC50, vin EC50 or the double and triple drug combination s, versus control (**Fig. S3, S4, S5, S6, Table S1)**. Protein enrichment analysis showed that DPI and MitoQ single treatments induced a similar profile of downregulated peptides, affecting mainly members of the complex I of the mitochondrial respiratory chain and cell cycle/division proteins (**Fig. S3, S4, Table S1**). However, they exerted contrasting effects on proteins involved in histone H3-K27 and H3-K4 trimethylation, with DPI causing downregulation and MitoQ causing upregulation. MitoQ upregulated the expression of proteins involved in cellular amino acid synthesis, and hydrogen peroxide catabolic processes. On the other hand, DPI mainly upregulated members of the cholesterol biosynthetic process (**Fig. S4, Table S1**). At its EC50 concentration, vincristine induced significant downregulation of Ribonucleotide Reductase M2 (RRM2), involved in DNA damage repair and cell cycle regulation (**Table S1)**. In agreement with their single-drug effects, both (M+D)/2 and (M+D+V)/3 downregulated proteins linked to mitochondrial ATP synthesis and cell cycle (**Fig. S5, S6, Table S1**). (M+D)/2 upregulated TTC5 and HMGCR, involved in p53-dependent apoptosis and cholesterol metabolic process, respectively (**Table S1**). The triple combination upregulated UBE2C, TTC5, ELOVL5 and NDUFS8. These proteins are implicated in mitosis, fatty acid metabolism, p53-dependent apoptosis and complex I of the mitochondrial respiratory chain, respectively (**Table S1**). Interestingly, the DNA damage repair protein RRM2, often upregulated in NB to support oncogenicity, was upregulated across all conditions, and phosphorylation at the Ser-80 and Ser-86 sites was significantly downregulated by both combinations (**Table S1**)^48–50^.

Based on the phosphoproteomic profiling we then inferred the upstream kinases responsible for the differential phosphorylation observed in each treatment versus control, using a kinase enrichment analysis (**Fig. 6**). Biological functions highly represented by the inferred kinases for both MitoQ and DPI single treatments include cell cycle regulation, DNA damage response and cell proliferation (DNAPK, CDK1, CDK2, RSK2, P7056K, UL97) (**Fig. 6A, B**). UL97 is a viral protein kinase that shares substrates with CDK1^51^. DNA damage response and regulation of cytoskeleton organization were also prominently represented in MitoQ EC50 treatment compared to control (LIMK1, LIMK2, TESK1, CAMK4, CAMK2B, PKACA, ERK1) (**Fig. 6B**). Moreover, MitoQ decreased MYCN phosphorylation at the Ser-156 residue (**Table S2**). No upstream kinases were predicted for vincristine on its own. For the double and triple combinations, the abundance in phosphorylation marks suggested that mainly kinases linked to cell cycle and cell proliferation are differentially regulated (UL97, PRKD1, CDK2, CDK4) (**Fig. 6C, D**).

**Fig 6.**
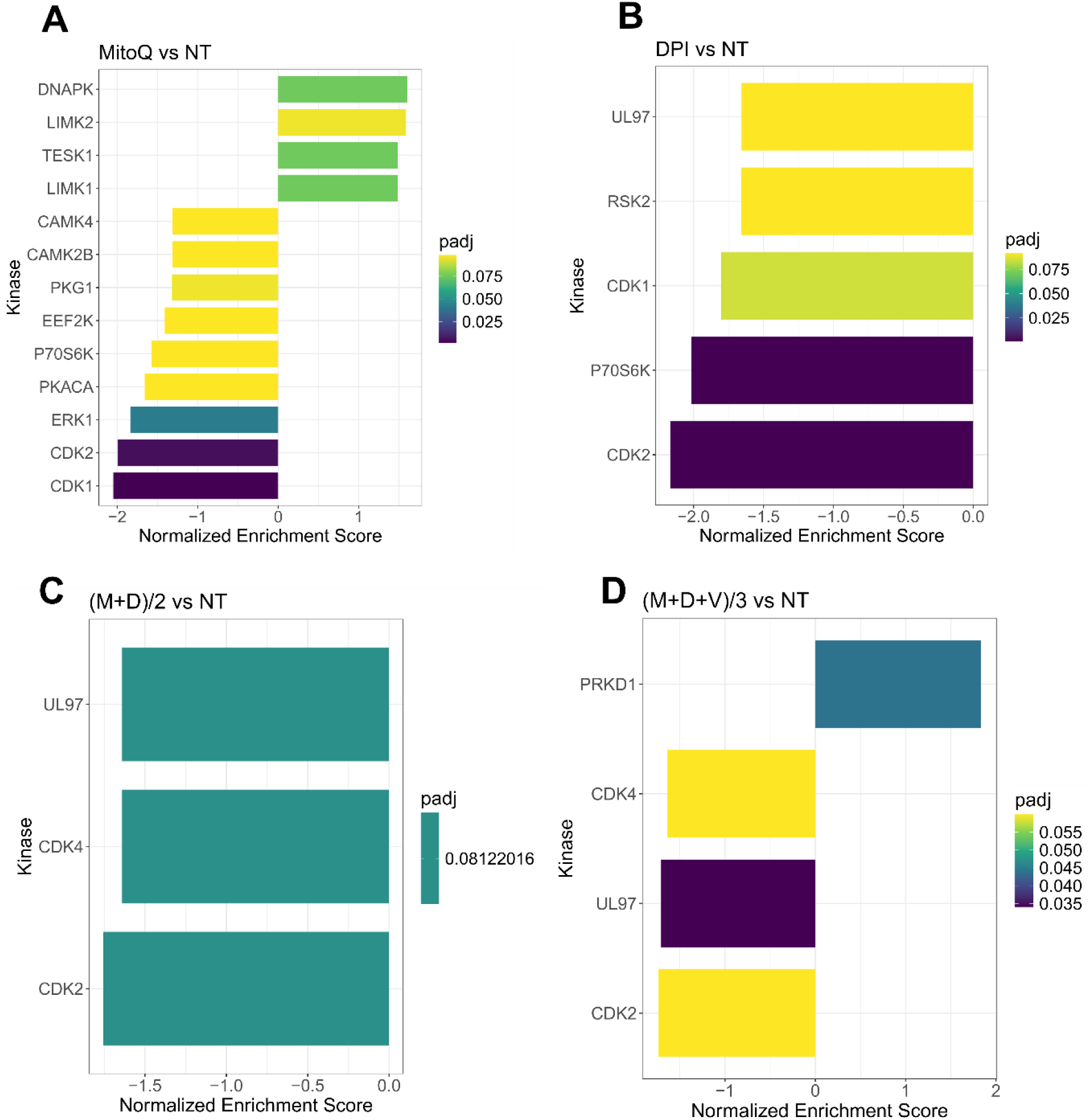
Kinase enrichment analysis of single and combined drug effects. Kinase enrichment inferred from phosphoproteomic data from **A)** MitoQ EC50 vs control, and **B)** DPI EC50 vs control, **C)** (M+D)/2 vs control, and **D)** (M+D+V)/3 vs control, after 24 hours of treatment. (Mean ± SEM; n=3; FDR = 0.05, s0 = 0.1).

Together, proteomic and phosphoproteomic profiling highlighted cell cycle and proliferation pathways as key targets of the double and triple drug combinations **(Fig. 6C, D, S5, S6)**. MitoQ, DPI and the double drug combination decreased the expression of cyclin D1 (G1/S-phase) and cyclin B1 (G2/M phase), suggesting an inhibitory effect on cell cycle progression. These findings were confirmed by Western blotting **(Fig. S7A)**. Similarly, the (M+D+V)/3 combination decreased the expression of cyclin B1 and, based on Western blot data, also downregulated cyclin D1. Vincristine caused a sharp increase in histone 3 phosphorylation at serine 10 (pS10-H3), likely due to its interference with mitotic spindle assembly, causing mitotic arrest (**Fig. S7A**). Since pS10-H3 is a marker for chromosome condensation linked to mitosis, cellular stress^52^ and DNA damage^53^, its elevation in the triple drug combination likely reflects both a proliferation block and a stress response to vincristine.

According to the MS results, MitoQ and to a lesser extent DPI reduced MYCN protein levels. These findings were confirmed by Western blotting (**Fig.S7**). However, this effect was reversed in Be(2)-C cells treated with the double and triple combinations (**Fig. S7A**). In contrast, in the IMR-5/75 MNA cell line, (M+D)/2 reduced MYCN by half, while (M+D+V)/3 led to an even greater decrease (**Fig. S7B**). This discrepancy may be related to the finding that Be(2)-C cells are more resistant to treatment and undergo apoptosis at a later time point.

Overall, these results suggest that the effects of MitoQ, DPI, and vincristine are mediated mainly through OXPHOS inhibition, MYCN regulation, and disruptions in DNA damage response and cell cycle progression.

To identify the proteins and pathways modulated by the combination treatments versus the single treatments, we applied a linear regression model to the matched global proteomic and phosphoproteomic data (**Fig. 7**). The model identifies pathways responsible for the effect of the double and triple drug combinations, versus the effect of the addition of each single agent at its EC50 concentration. The differentially expressed proteins revealed that mainly the respirasome, mitochondrial ribosome, hydrogen peroxide and glutathione metabolic processes are responsible for the effect exerted by the double combination on Be(2)C cells (**Fig. 7A**). Regarding the triple combination, the pathways are highly similar to those of (M+D)/2, with an additional enrichment in pathways associated with anion transmembrane transporter activity (**Fig. 7B**). In terms of kinase enrichment, pathways related to cell cycle, DNA damage response, cell proliferation and regulation of cytoskeleton organization account for the effects of (M+D)/2 and (M+D+V)/3 compared to the addition of single agents (LIMK1, TESK1, P70S6K, CDK1, CDK2, CAMK4, CAMK2B) (**Fig. 7C, D**). These findings suggest that the mechanism of synergy underlying the interaction between DPI and MitoQ is linked to their impact on mitochondrial metabolism and cell cycle regulation. Furthermore, the addition of vincristine does not significantly alter the proteomic profile linked to the activity of the other two compounds, while still causing more cell death when combined with MitoQ and DPI at a low dosage. The independent function of vincristine underscores the therapeutic potential for MitoQ and DPI to engage previously untargeted pathways involved in cancer progression.

**Fig 7.**
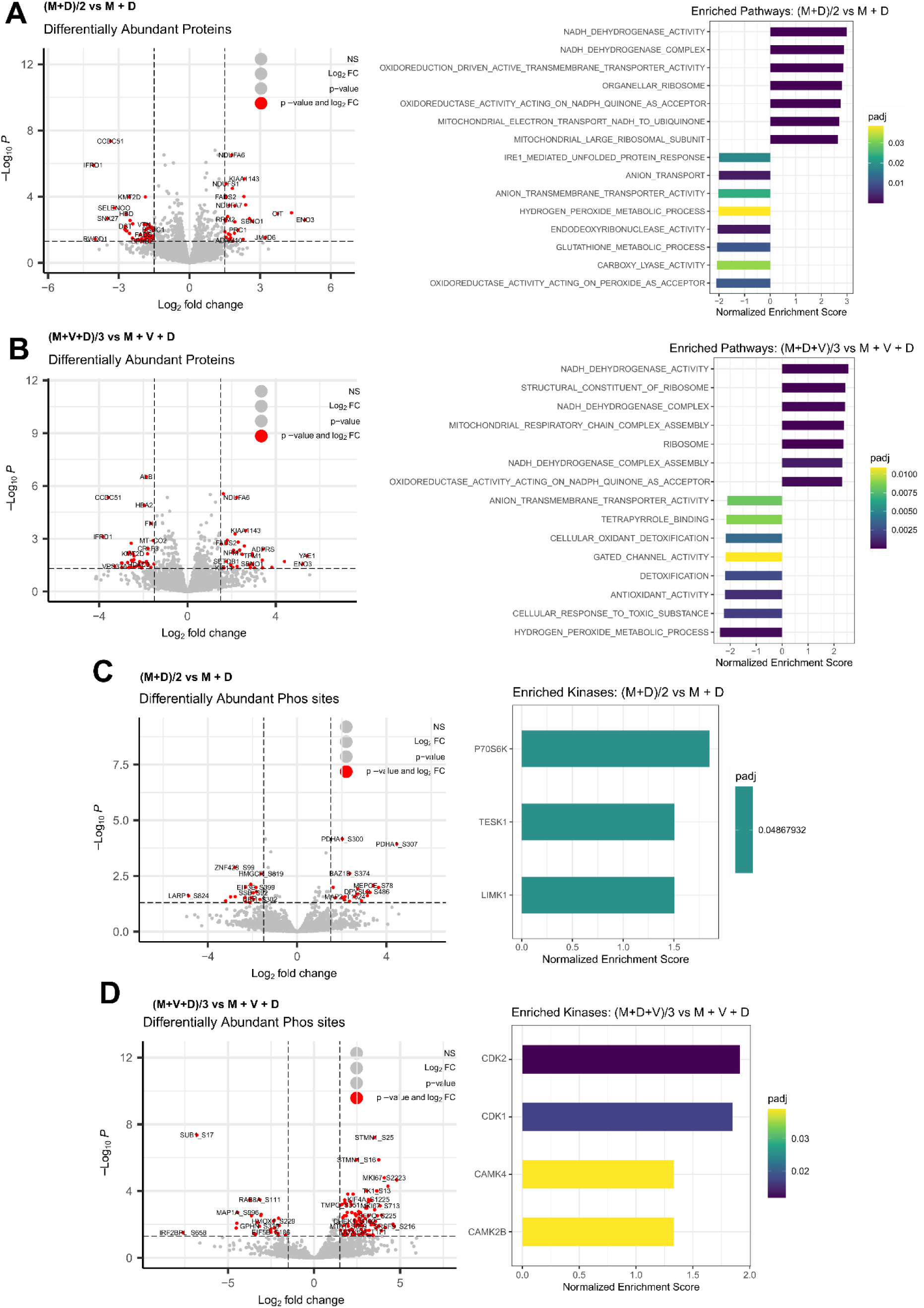
Integrated phosphoproteomic and proteomic analysis of drug synergies. Differentially enriched proteins and associated pathways following treatment of Be(2)-C MNA cells for 24 hours with **A)** (M+D)/2 vs MitoQ EC50 + DPI EC50, or **B)** (M+D+V)/3 vs MitoQ EC50 + DPI EC50 + Vin EC50. (Mean ± SEM; n=3; adj p value < 0.05, s0 = 0.1). Inferred kinases from phosphoproteomic data following 24 hours treatment of Be(2)-C cells with **C)** (M+D)/2 vs MitoQ EC50 + DPI EC50, or **D)** (M+D+V)/3 vs MitoQ EC50 + DPI EC50 + Vin EC50. (Mean ± SEM; n=3; adj p value < 0.05, s0 = 0.1)..

### Potential molecular mechanisms of drug synergies

In both the double and triple drug combinations, processes generating ROS (NADH dehydrogenase, NADH quinone oxidoreductase/complex I) are significantly enriched, while antioxidant activities are significantly decreased (**Fig. 7A, B**). Therefore, we examined superoxide and H_2_O_2_ generation in MNA cells (**Fig. 8**). Specifically, we exposed Be(2)-C cells to conditions akin to those used for MS analysis, utilizing either DPI, MitoQ and vincristine individually or in combination. Indeed, superoxide production has previously been demonstrated to be linked to the susceptibility of MNA NB cells to DPI, resulting in cytochrome c release, caspase activation, cytochrome c release and cell death^31^. MitoSOX red dye (Invitrogen) was used as a probe to monitor mitochondrial superoxide, and Abgreen indicator (Abcam) for H_2_O_2_. The Abgreen indicator (Abcam) is a cell-permeable fluorescent indicator which generates green fluorescence when reacting with H_2_O_2_. 10 μM DPI was used as a positive control for superoxide production, while 10 μM doxorubicin was used as a positive control for H_2_O_2_ generation. DPI, MitoQ, and vincristine at their EC50 concentration s individually increased mitochondrial superoxides over a 24-hour period (**Fig. 8A**). Interestingly, MitoQ decreased H_2_O_2_ after 2 hours of treatment, but this effect was reversed after 24 hours (**Fig. 8B**). Both (M+D)/2 and (M+D+V)/3 combinations increased superoxide production after 24 hours, as well as H_2_O_2_ (**Fig. 8A, B**). Thus, the increase in H_2_O_2_ production for the combinations after 24 hours reflects the enrichment analysis of the proteomics data, which indicate enhanced ROS production and less ROS removal (**Fig. 7B**). These results further suggest that a balanced generation of ROS by mitochondrial complex I is crucial to maintain MNA cellular fitness. Overall, these results confirm what was observed in the proteomic analysis and the mechanism of synergy.

**Fig 8.**
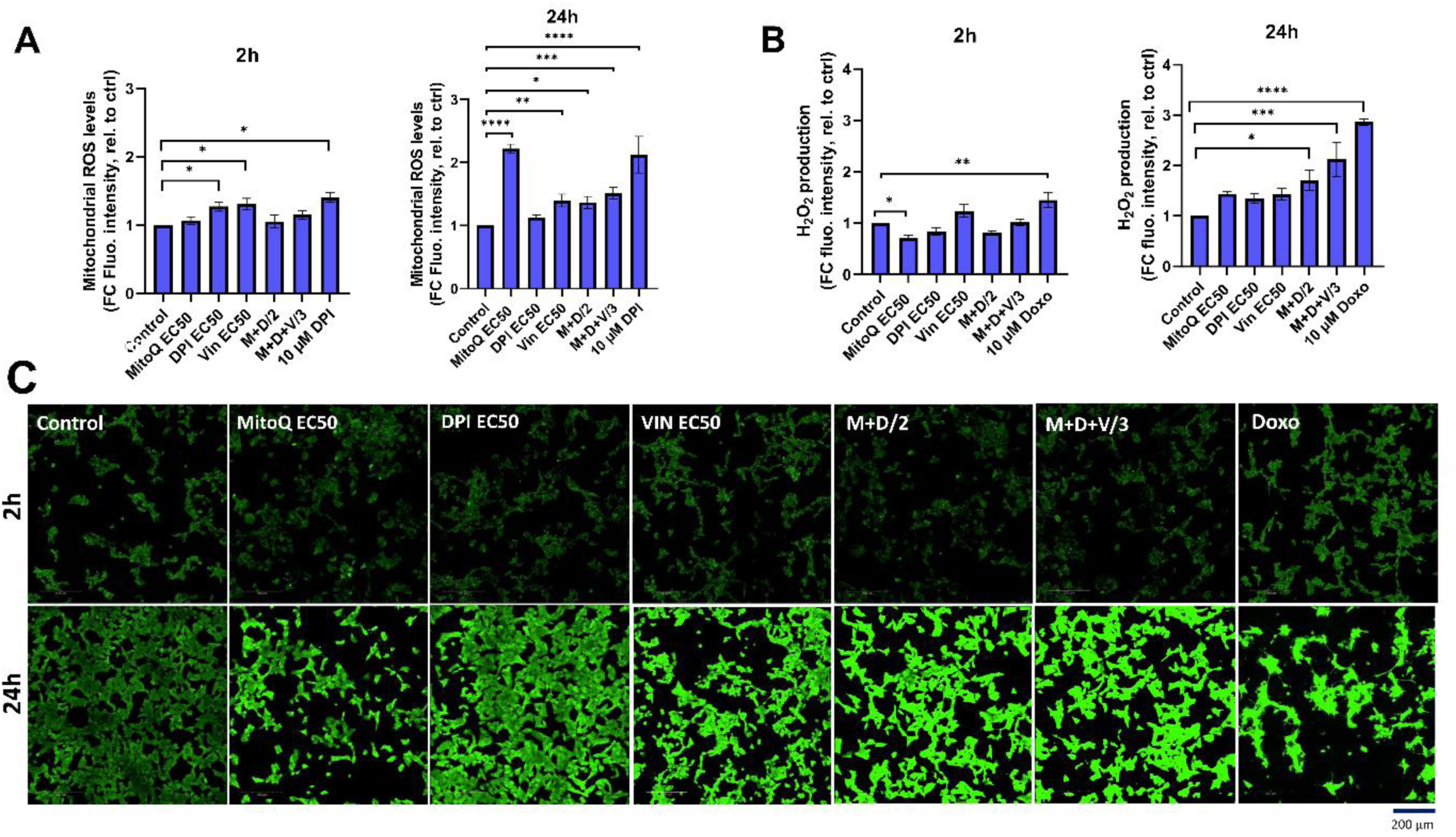
Potential molecular mechanisms of drug synergies. **A)** MitoSOX Red dye was used to measure mitochondrial superoxide production in Be(2)-C under the same treatment conditions as in the proteomic analysis, for 2h, and 24h. (Mean ± SEM; n=3; Ordinary One-Way ANOVA, p<0.05 = *, p<0.005 = **, p<0.0005 = ***, p<0.0001 = ****). 10 μM DPI was used as a positive control for increase in mitochondrial superoxides. Scale bar = 200 μM. **B, C)** AbGreen indicator dye was used to measure H2O2 production in Be(2)-C under the same treatment conditions as in the proteomic analysis. Doxorubicin was used as a positive control for H2O2 induction. (Mean ± SEM; n=3; Ordinary One-Way ANOVA, p<0.05 = *, p<0.005 = **, p<0.0001 = ***). Below, associated confocal images x40. Scale bar = 200 μM.

### *In vivo* evaluation of MitoQ and DPI: Tolerability and limited efficacy in NB mouse models

Finally, to evaluate the efficacy of MitoQ and DPI *in vivo*, both drugs were administered individually and in combination to Th-*MYCN* transgenic NB mice. Despite the promising efficacy observed in Th-*MYCN*-derived 3D spheres, administration of the compounds resulted in gastrointestinal toxicity, including blocked intestine and stomach enlargement, necessitating early termination of the experiment. To address this limitation, the study was re-conducted using an IMR-5 (MNA) xenograft immunodeficient NB mouse model, where a reduced toxicity was observed (**Fig. 9**). In contrast to our *in vitro* results, treatment with MitoQ, DPI, or their combination did not produce notable effects on tumor size reduction, even at the highest tolerable doses (**Fig. 9A**). While a trend toward increased survival probability was observed in animals treated with either compound or their combination, the effect was not statistically significant (**Fig. 9B**). These findings suggest that MitoQ and DPI are more tolerated in an immunodeficient *in vivo* environment, but their limited efficacy in reducing tumor burden may arise from challenges linked to oral administration, including poor absorption and first-pass metabolism. Alternatively, these findings may point to a role of the immune system in the effect of these drugs *in vivo*. We are currently investigating possible reasons for this discrepancy.

**Fig 9.**
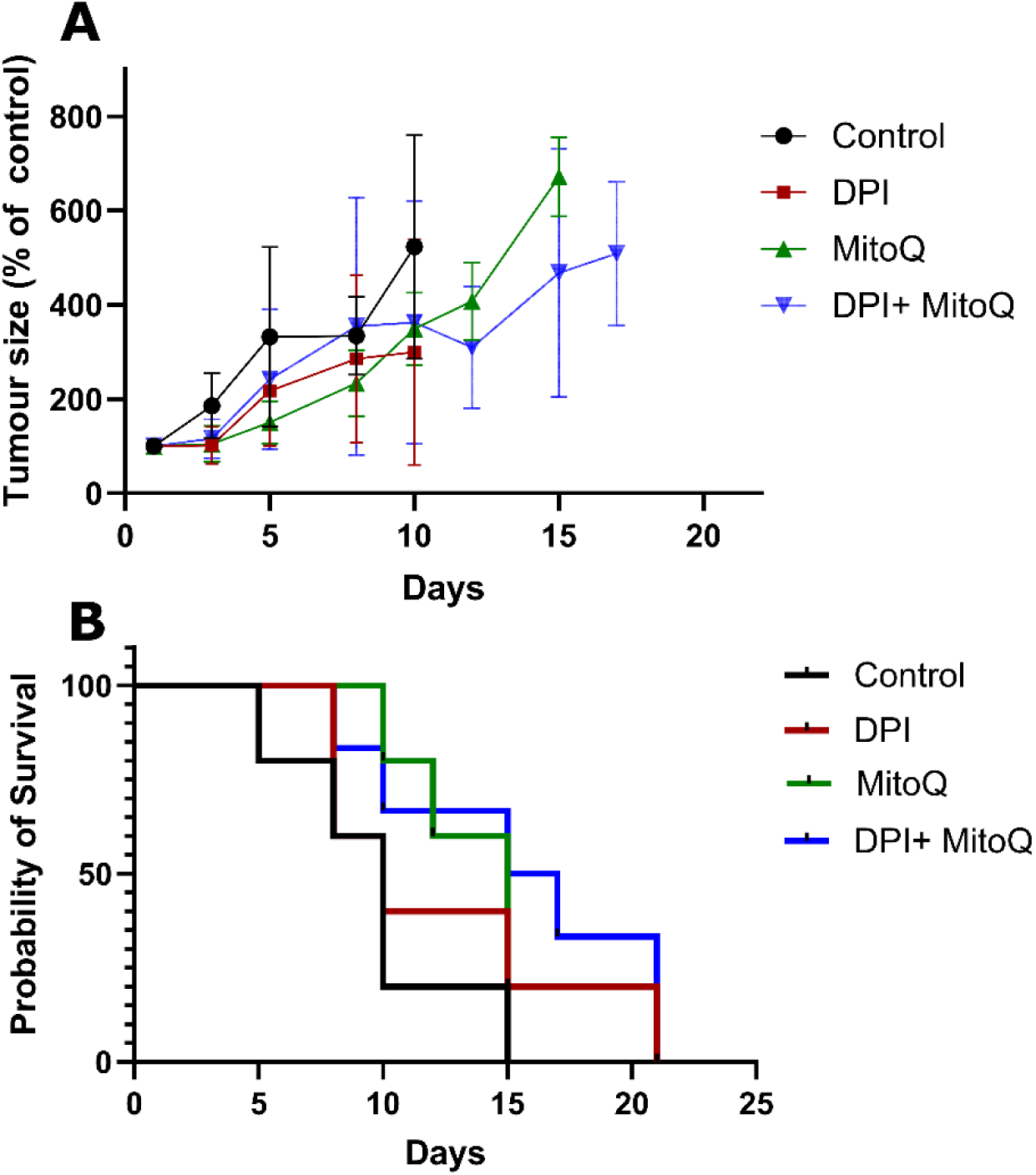
In vivo treatment of the IMR-5 xenograft immunodeficient NB mouse model. **A)** Tumor size normalized to untreated control, measured over a 20-day treatment course with 3 mg/kg DPI, 30 mg/kg MitoQ, or their combination (1.5 mg/kg DPI and 20 mg/kg MitoQ). Efficacy curves terminate when 50% of the animals in a treatment arm had died. **B)** Probability of survival for each treatment group, calculated based on survival events. Each treatment group consisted of at least 5 mice. Drugs were administered via oral gavage every two days. The endpoint of the trial was reached when the tumors grew to the size limit (15 mm in diameter). (Mean ± SEM). Log-rank (Mantel-Cox) test p=0.2; Logrank test for trend p=0.0482; Gehan-Breslow-Wilcoxon test p=0.2087.

## 3. Discussion

Targeting MYCN directly in NB has been challenging, primarily due to the absence of drug-binding pockets in MYCN. Using combination therapies to target critical downstream signaling pathways is considered a rational approach.

In this paper, we initially focused on uncovering the mechanism of action of DPI, a NADPH oxidase inhibitor that we previously found to specifically inhibit MNA NB cell proliferation^31^. DPI was found to increase mitochondrial superoxide production in MNA cells, leading to the induction of an apoptotic cascade^31^. As a result, MitoQ was chosen as a mitochondria-specific ROS inhibitor to decrease superoxides and assess if it could reverse DPI-induced cell death on MNA cells. MitoQ is derived from the lipophilic molecule ubiquinone, allowing it to efficiently pass through lipid bilayers and accumulate within the inner mitochondrial membrane of cells^54,55^. Inside the mitochondrion, MitoQ undergoes redox cycling mediated by complex II, where it is reduced to mitoquinol, known for its potent antioxidant properties^56^. Notably, this compound exhibits protective effects on normal cells^57^ and is being investigated as a potential therapeutic agent for neurodegenerative and aging-related conditions^58,59^. Surprisingly, we found that MitoQ alone increased mitochondrial superoxide production. While it reduced high-dose DPI-mediated ROS, it did not prevent cell death of MNA NB cells. MitoQ exacerbated DPI-induced cytotoxicity and was found toxic on its own at low nM concentrations. Consistent with these findings, other groups have shown that MitoQ can enhance superoxide production ^60^, and cause cytotoxicity in tumor cells^32,61^. Moreover, MitoQ appears to exhibit a MYCN-dependent inhibitory effect on MNA NB cells similar to that of DPI^16^. When administered alone, MitoQ increased neuronal differentiation of MNA NB cells, suggesting a transition towards a more “favorable” phenotype. Additionally, we observed strong synergism between DPI and MitoQ in reducing cell viability of MNA cells and a 3D MYCN-driven NB tumor model. The efficacy of the combination in a spheroid model of NB provides confidence that it may translate to *in vivo* models. These findings strongly suggest that MitoQ enhances the efficacy of DPI while also exhibiting antitumor activity as a standalone treatment in NB.

Kong et al., similarly uncovered synergy between a complex I inhibitor and a mito-antioxidant in cancer cells, suggesting their ability to reduce mitochondria-derived H_2_O_2_ levels as a potential mechanism of action^28^. In our study, H_2_O_2_ levels increased after 24 hours of (M+D)/2 and (M+D+V)/3 treatments. We, therefore, believe that the observed cytotoxic effect may not be directly mediated by a reduction in H_2_O_2_ in this context, but rather the opposite. MitoQ and DPI-induced damage to the respiratory chain may increase the NADH/NAD+ ratio over time, leading to O_2_ leakage and subsequent superoxide formation ^62^. Indeed, high reductive stress can paradoxically elevate superoxide levels via Glutathione Reductase (GSR) and Thioredoxin Reductase (TrxR) due to impaired electron flow caused by inhibitors^63^. In line with these findings, we previously found that high-dose DPI (10 μM) increased Glutathione Peroxidase 1 (GPX1), GSR and Glutathione Synthetase (GSS) protein expression levels in MNA cells^31^ (**Fig. S8**). Our linear regression model also highlighted antioxidant response and glutathione metabolic processes as key pathways responsible for the activity of MitoQ + DPI, which could be the result of a high H_2_O_2_ production. Furthermore, it is crucial to note that superoxides are the primary but not the sole source of H_2_O_2_. H_2_O_2_ is also generated via the peroxisome, endoplasmic reticulum, and certain NOX enzymes, such as NOX4^64^. Although it is difficult to conclude that ROS are the primary cause of apoptosis in this context, it is more likely that both DPI and MitoQ cause irreversible damage to OXPHOS in MNA cells^23,32^. Recent studies also demonstrated that redox-crippled MitoQ inhibited the proliferation and metastasis of breast cancer and glioma cells, suggesting that this effect is due to the inhibition of complex I rather than antioxidant properties^32,65^. “Redox-crippled” MitoQ is a modified version of the mitochondrial-targeted antioxidant with its redox activity removed, allowing differentiation between effects from its antioxidant properties and other factors^32^.

Adding vincristine to the MitoQ-DPI combination was motivated by its common use in the conventional clinical treatment of high-risk NB^66^. Reducing the use and concentration of chemotherapeutic agents through combination therapy represents a key step towards less toxic treatments. We found that vincristine and DPI rather had an additive than a synergistic interaction. While synergism and additivity allow for achieving better effects with lower drug concentrations, clinical data suggest that independent drug action can also significantly improve patient outcomes^67^. One way of assessing drug synergy consists of combining drugs at half of their EC50 concentration for a two-drug combination, and one-third of their EC50 concentration for a three-drug combination^40,68^. Both (M+D)/2 and (M+D+V)/3 synergistically reduced cell viability in two MNA cell lines. These combinations also increased cell death and induced caspases 3/7 activation in all tested high-risk NB cells.

The use of MYCN-regulatable cell systems only moderately affected treatment sensitivity, likely due to an incomplete regulation of MYCN protein expression levels. MYCN downregulation in MNA IMR-5/75 cells only achieved 50% reduction in MYCN levels, which is a higher expression than in MYCN non-amplified NB. Similarly, SH-SY5Y/MYCN +Dox increases MYCN overexpression, but these levels remain significantly lower than those seen with MNA. Interestingly, similar to DPI, MitoQ alone potently reduced MYCN levels in our most resistant NB cell line, Be(2)-C. Although the double and triple combinations at reduced doses did not decrease MYCN levels after 24 hours, they potently diminished MYCN protein levels in the IMR-5/75 cell line. Moreover, MitoQ was found to decrease MYCN phosphorylation at the Ser-156 residue. This phosphorylation site resides within a disordered region lacking a defined structure; thus, its functional implications remain unclear. The reduction of MYCN protein levels in MYCN-driven brain tumors is of major importance due to its central role as an oncogenic driver^69^.

In addition to mitochondrial damage, whole proteomic and phosphoproteomic analysis highlighted the impact of our drug combinations on cell cycle blockade and DNA damage as part of their mechanism of action. Notably, vincristine showed minimal influence on the proteomic landscape altered by MitoQ + DPI combination. Interestingly, we consistently observed downregulation of RRM2 at the protein level across all single and combination treatments. Moreover, both (M+D)/2 and (M+D+V)/3 significantly decreased phosphorylation of RRM2 at Ser-80 and Ser-86 residues. RRM2 is an enzyme playing a crucial role in DNA synthesis and repair via deoxyribonucleotide production ^50^. Importantly, RRM2 is often found overexpressed in NB, and its depletion was shown to induce G1/S phase arrest due to replicative stress and checkpoint kinase 1 (CHK1) activation ^48,49^. Elevated RRM2 levels also correlate with MYCN expression levels and poorer survival outcomes in NB patients^48^. Although CHKs did not show differential expression at 24 hours post-treatment, it is plausible that such changes may manifest at later timepoints subsequent to RRM2 downregulation. (M+D)/2 and (M+D+V)/3 also upregulated the protein expression of mitochondrial Tetratricopeptide Repeat Domain 5 (TTC5/STRAP). TTC5 has previously been shown to downregulate ATP synthase activity, which sensitizes cells to mitochondrial-dependent apoptosis via enhanced p53 activity^70^. This protein exerts a dual role, as it also accumulates in the nucleus following DNA damage, thereby enhancing the activity of p53 and leading to cell cycle and apoptosis. Moreover, TTC5 is a MYC interactor in acute myeloid leukemia stem cells, preventing excessive accumulation of MYC^71^. While there is no direct evidence of TTC5 interaction with MYCN in NB, both proteins are direct interactors of the histone acetyltransferase _p30072,73._

Therefore, we propose that both combinations induce cytotoxicity through two potential mechanisms. The first one involves irreversible damage to the ETC, possibly through TTC5 upregulation, resulting in ATP loss, increased mitochondrial superoxide production and caspases 3/7 activation, and subsequent apoptosis. The second mechanism involves inhibition of clonogenic cell growth, potentially via downregulation of RRM2 and DNA damage, leading to accumulation of DNA breaks and cell cycle blockade. Microscopy experiments on a 3D spheroid model of MYCN-driven NB support this hypothesis. Indeed, an increase in DNA fragmentation and condensation was visible one day after the addition of M+D and M+D+V.

Several complex I inhibitors have been evaluated clinically and discontinued due to neurotoxicity and adverse side effects (IACS-010759, Tamoxifen, Piericidin A, Phenformin)^74,75^. MitoQ was found to induce mitochondrial damage in mouse kidney cortex tissue^76^, although the concentrations used were much higher than the typical dosage used for cancer cells. However, over 400 peer-reviewed studies have reported on the therapeutic benefits of MitoQ, demonstrating its protective effects against a range of pathologies^77^. Furthermore, MitoQ has been evaluated in several clinical trials and has shown overall good tolerability, with nausea and vomiting being the most commonly reported side effects^77^. Additionally, MitoQ exhibits cardioprotective effects in rodents during doxorubicin-induced cardiac damage^78^, or induced-sepsis^79^. The major challenge seems to be establishing a standardized dosage of MitoQ for specific human diseases. Our results suggest that MitoQ can be used at low nanomolar concentrations across various MNA NB cell lines, while having a MYCN-dependent effect. In regard to DPI, we previously demonstrated that low micromolar concentrations effectively reduced tumor growth and metastasis in a MYCN-driven zebrafish model of NB, with low associated toxicity^31^. The MYCN-driven effects of both DPI and MitoQ can potentially limit off-target effects and minimize damage to healthy cells.

A limitation of this study is the discrepancy between the strong *in vitro* synergy of the MitoQ + DPI combination and its limited effectiveness in *in vivo* models of NB. The *in vivo* environment is considerably more complex, involving intricate biological networks and an active immune system, which are not present in in vitro cultures. Furthermore, gastrointestinal toxicity observed in the transgenic mice suggests that bioavailability may be compromised with oral administration. This can hinder drug effectiveness, as some drugs degrade in the acidic stomach environment or due to gut enzymes^80^. While intravenous or intraperitoneal routes may improve bioavailability, they are difficult to apply in small animal models. The observed differences appear to be dependent on the animal model, as previous studies have shown that MitoQ is well tolerated at concentrations up to 500 μM in drinking water over several weeks, without causing adverse effects on behavior or metabolism^55,81^.

In conclusion, our data provides therapeutic opportunities for targeting MNA NB growth. The present study reveals that two mitochondria targeted agents, MitoQ and DPI were highly synergistic in reducing cell viability of MNA NB cells and MYCN-driven NB spheroids. MitoQ exhibited great antitumor activity as a monotherapy against MNA cells, showing a MYCN-dependent effect similar to DPI. Moreover, we showed that MitoQ greatly decreased MYCN protein levels in MNA cells. The combination of MitoQ and DPI with a widely used chemotherapeutic agent allowed to more closely resemble potential clinical treatments for NB patients. Moreover, reducing drug dosage still achieved significant cytotoxic effects. Our data suggest that the tested drug combinations induce cell death in MNA cells through two mechanisms: inhibition of OXPHOS and cell cycle arrest. A common concern in drug synergy is the focus on combinations of agents with limited to null monotherapy activity^82^. We demonstrate here that our single agents are highly effective at low concentrations on their own and show substantial synergistic effects when combined.

## 4. Materials and Methods

### 4.1. Cell Culture and *in vivo* experiments

#### Cell Lines

Four human NB cell lines were used in this paper: Be(2)-C (MNA), IMR5-/75-shMYCN^29^ (MNA, doxycycline-inducible shRNA targeting MYCN, resulting in MYCN downregulation), IMR-32 (MNA), and SH-SY5Y/6TR(EU)/pTrex-Dest-30/MYCN (non-MNA, doxycycline-inducible transgene, resulting in MYCN overexpression). IMR-5/75-shMYCN and SH-SY5Y/6TR(EU)/pTrex-Dest-30/MYCN are named IMR-5/75 and SH-SY5Y/MYCN, respectively, in this paper. The cell lines were generously provided by Dr. Johannes Schulte and Dr. Frank Westermann. Cell lines were grown under standard tissue culture conditions in an incubator (37 °C and 5% CO_2_). All cell lines were cultured in RPMI 1640 media (Gibco) supplemented with 10% Fetal Bovine Serum (FBS) (Gibco), 1% L-Glutamine (Gibco) and 1% Penicillin/Streptomycin (Gibco). For IMR-5/75, the culture media was supplemented with 5 μg/mL Blasticidin (Bio-Sciences) and 50 μg/mL Zeocin (Thermo Scientific) for plasmid selection. For SH-SY5Y/MYCN, the culture media was supplemented with 7.5 μg/mL Blasticidin (Bio-Sciences) and 200 μg/mL Geneticin (G418) (Thermo Scientific) for selection of the plasmid. 1 μg/mL doxycycline (Sigma-Aldrich) was added to the media for 24 h to induce the expression of an shRNA against MYCN in IMR-5/75 cells and overexpress MYCN in SH-SY5Y/MYCN cells.

#### 3D Spheroid NB Model

The primary murine NB neurospheres were generously provided by Dr. Evon Poon and Prof. Louis Chesler (ICR, England). These cells were extracted from tumors coming from Th-*MYCN* transgenic mice over-expressing *MYCN* gene under the control of the tyrosine hydroxylase promoter. Spheroids were grown under standard tissue culture conditions in an incubator (37 °C and 5% CO_2_). The cells were cultured in Ultra-Low-Attachment plates (ULA) (Thermo Scientific) in PrimNeuS media^83^.

#### Mice and *in vivo* experiments

All animal experiments were approved by The Institute of Cancer Research Animal Welfare and Ethical Review Body and performed in accordance with the UK Home Office Animals (Scientific Procedures) Act 1986, the UK National Cancer Research Institute guidelines for the welfare of animals in cancer research and the ARRIVE (animal research: reporting in-vivo experiments) guidelines. Female NSG mice were obtained from Charles River and enrolled into trial at 6–8 weeks of age. Mice were maintained on a regular diet in a pathogen-free facility on a 12 h light/dark cycle with unlimited access to food and water.

One million IMR5 cells with 50% matrigel were injected subcutaneously into the right flank of NSG mice (female; 6–8 weeks old) and allowed to establish a murine xenograft model. Studies were terminated when the mean diameter of the tumor reached 15mm. Mice bearing IMR5 xenografts with a mean diameter of 5 mm were treated with 3 mg/kg DPI (P.O., 5 days/week), 30 mg/kg MitoQ (P.O., 3 days/week), or their combination (1.5 mg/kg DPI (P.O., 5 days/week) and 20 mg/kg MitoQ (P.O., 3 days/week)). Tumor volumes were measured by Vernier caliper across 2 perpendicular diameters, and volumes were calculated according to the following formula: V = 4/3π [(d1 + d2)/4]3 where d1 and d2 are the 2 perpendicular diameters. The endpoint of the trial was reached when the tumors grew to the size limit (15 mm in diameter).

### 4.2. Chemicals

Mitoquinone Mesylate (MitoQ, MQ) was purchased from MedChemExpress (#HY-100116A), and Diphenyleneiodonium chloride (DPI) from Enzo Life Sciences (#BML-CN240-0010). Dimethyl sulfoxide (DMSO, vehicle) (#D2650), Vincristine Sulfate (Vin) (#V8388) and retinoic acid (#R2625) were from Merck. Doxorubicin was purchased from Selleck Chemicals (#S1225).

### 4.3. CellTox Green Assay

Cell toxicity was evaluated using the CellTox Green fluorescent assay (Promega). Typically, cells were seeded in 50 μL media per well in duplicates on a black 96 -well plate with clear bottom (Thermo Scientific). Complete RPMI phenol red-free media (Gibco) was used and was beforehand completed with CellTox Green Dye at a 1:500 ratio (v/v). Six h after seeding (or 24 h after doxycycline induction for MYCN-regulatable cell systems), the cells were treated with 50 μL of drug(s) or vehicle and incubated at 37 °C for 72 h. Fluorescence was measured at 4, 24, 48 and 72 h after treatment at 485 nm excitation and 530 nm emission on a SpectraMax M3 microplate reader (Molecular Devices).

### 4.4. MitoSOX Red and PI Stainings

Mitochondrial superoxide was measured with MitoSOX Red fluorescent indicator (Thermo Scientific). Propidum iodide (PI) (Thermo Scientific) was used to selectively stain apoptotic cells. Typically, cells were seeded in 100 μL RPMI phenol red free media per well in duplicates on a black 96-well plate with clear bottom (Thermo Scientific). To prevent IMR-5/75 from detaching, the plates were treated beforehand with 100 μg/mL Poly-D-Lysine (Sigma-Aldrich). Six h after seeding (or 24 h after doxycycline induction for IMR-5/75), the cells were treated with 100 μL of drug(s) or vehicle and incubated at 37 °C. After the appropriate incubation time, the cells were washed once with DPBS Ca^2+^/Mg^2+^ (Invitrogen). MitoSOX Red dye (4 μM) or PI (1 μg/mL) were added together with Hoechst nuclear counterstain (1 μg/mL) (Bio-Sciences) to the cells in DPBS Ca^2+^/Mg^2+^ and incubated for 20 minutes at 37 °C protected from light. The cells were washed twice with DPBS Ca^2+^/Mg^2+^, and fresh RPMI phenol red-free media was added before live cell imaging. MitoSOX Red indicator and PI were detected using the TRITC filter (Ex/Em: 396/610 nm) and Texas Red (Ex/Em:560/630), respectively, while Hoechst was detected using HOECHST 33342 filter (Ex/Em: 353/483).

### 4.5. Live Cell Imaging

Confocal images of 4 optical planes interspaced by 1 μm (from -1.0 μm to 2.0 μm included) across 25 x 25 fields of view with a 10% overlap were acquired per well on an Opera Phenix High Content Screening System (PerkinElmer) using a 40x or 63x/1.15 NA water-immersion objective, with live cell parameters (37 °C and 5% CO_2_). The acquired images were analyzed using an optimized pipeline for each cell line on Harmony® v.4.9 high-content analysis software (PerkinElmer). The building blocks within the software were used to identify and calculate the mean dye parameters per well: number of nuclei, MitoSOX Red, or PI mean intensity/cell.

### 4.6. Neuronal Differentiation

IMR-5/75 cells were plated in duplicates on a 6-well plate in 2 mL complete RPMI phenol red-free media (Gibco). One day after seeding, the cells were treated with either 50 nM MitoQ, 100 nM MitoQ, 10 μM retinoic acid or vehicle, and incubated 37 °C for 7 days. Every two days, the cells were treated with the appropriate drug or vehicle, in 2 mL fresh media. After 7 days, the cells were fixed, permeabilized and neuronal differentiation was assessed by Alexa Fluor 488 Phalloidin staining (Bio-Sciences) according to the manufacturer’s instructions. Hoechst nuclear counterstain (1 μg/mL) (Bio-Sciences) was used for nuclei counting. Imaging was performed on an EVOS Digital Inverted Fluorescence Microscope (Invitrogen), using the FITC (green) and the DAPI (blue) filters. Neurite outgrowth (cells with neurites >= 2 cell body diameters) were counted per cell using the NeuriteJ plugin of ImageJ (v1.53e; Java 1.8.0_172).

### 4.7. CellTiter-Glo Assay

Cell viability by loss in ATP content was measured using the CellTiter-Glo Luminescent and CellTiter-Glo 3D Luminescent Cell Viability Assays (Promega). Cells were seeded in duplicates in 100 μL media on a Nunc 96-Well White Microplate (Fisher Scientific). Six h after seeding, the cells were treated with 100 μL of drug(s) or vehicle and incubated at 37 °C for 72 h. To assess cell viability of the neurospheres, cells were seeded after dissociation in 100 μL PrimNeuS media on a Nunclon Sphera 96-Well U-Shaped-Bottom Microplate (Thermo Scientific). Two days after seeding, the cells were treated with 100 μL of drug(s) or vehicle and incubated at 37 °C for 72 h. CellTiter-Glo reagent was added to the wells according to the manufacturer’s instructions and luminescent signal was measured on a SpectraMax M3 microplate reader (Molecular Devices).

### 4.8. Western blotting

Six hours after seeding (or 24 h after doxycycline induction for IMR-5/75), the cells were treated with either MitoQ EC50, DPI EC50, Vin EC50, (M+D)/2, (M+D+V)/3 or vehicle and incubated at 37 °C for 24 h. Cells were harvested using 110 μL ice cold lysis buffer containing 20 mM Tris-HCl (pH 7.5), 150 mM NaCl, 1 mM MgCl_2_, 1% (v/v) Triton X-100 and 10% (v/v) of protease and phosphatase inhibitor cocktail (Roche). After sample denaturation and electrophoresis, membranes were blocked with 5% milk in TBS-T (120 mM Tris-HCl, pH 7.4, 150 mM NaCl, and 0.05% Tween 20) for 1 h at room temperature. Target proteins were detected using primary antibodies at 1:1000 dilution overnight at 4°C (MYCN #sc-53993, Santa-Cruz Technologies; Phospho-Histone H3 (Ser10) #3465, Cell Signaling Technology; Phospho-Histone H2AX (Ser139) #05-636, Sigma-Aldrich; Cyclin B1 #12231, Cell Signaling Technology; Cyclin D1 #2922, Cell Signaling Technology; GAPDH #2118, Cell Signaling Technology). Anti-mouse or anti-rabbit secondary antibodies (Cell Signaling Technology) HRP-conjugated at 1:5000 dilution were incubated with the membranes for 1 h at room temperature, and visualized using a chemiluminescence system. GAPDH was used as a loading control.

### 4.9. RT-qPCR

IMR-5/75 cells were seeded in 6-well plates and collected 24 h after doxycycline induction. RNA isolation was performed using the RNeasy Mini Kit (#74104, Qiagen) and cDNA synthesis using the High Capacity cDNA Reverse Transcription Kits with RNase Inhibitor (#4374966, Thermo Scientific) according to the manufacturer’s instructions. Polymerase chain reactions were performed using TaqMan Gene Expression assays (Bio-Sciences) for *MYCN* (#Hs00232074_m1) and two housekeeping genes, *GAPDH* (#Hs02786624_g1) and *B2M* (Hs00187842_m1) on a QuantStudio 7Flex qPCR System (Applied Biosystems). All reactions were performed in duplicates and the same amount of cDNA was used for all reactions. Ct Mean values of the two housekeeping genes (*GAPDH* and *B2M*) were averaged for each condition and these values were used for normalization. The average Ct Mean of both reference genes is then subtracted from the Ct value of *MYCN*, for each sample: ΔCt = Ct Mean(*MYCN*) − Ct Mean(*GAPDH/B2M*).

### 4.10. Synergy Analysis

Synergy scores were calculated using SynergyFinder 3.0 for 2-drug combinations. Normalized cell viability values were imported to SynergyFinder 3.0 in a matrix format, and %viability was chosen as a readout. LL4 was selected for curve-fitting algorithm, detection of outliers was activated, and the Zero Interaction Potency (ZIP) or Loewe model were used to capture the drug interaction relationships.

To investigate the potential synergy of two or more drugs in combination at reduced doses, the Bliss Independence model was used. According to Rukhlenko et al.^40^, the Bliss synergy score was defined as follows for reduced drug combinations:

According to the Bliss criterion, when two independent drugs are combined, their effects simply add up, and the Bliss synergy score is zero. In that case, if the combination is merely additive, the effect would be the following:

**E_expA+B_ = √E_A_*√E_B_**

In this equation, E is the %viability effect obtained from the CellTiter-Glo output. E_A+B_ is the observed effect of the combined drugs, and E_A_*E_B_ the expected effect when the drug effects are independent (viability values of the two single drugs). The Bliss synergy score was therefore calculated as follows:

**S_bliss_ = E_expA+B_ – E_obsA+B_ = √E_A_*√E_B_ - E_A+B_**

In this context, a positive Bliss score means that the observed effect of the combination is lower than what would be anticipated if the drug effects were independent, thereby indicating synergy. In contrast, a negative Bliss score would suggest antagonism.

### 4.11. Total Proteomics and Phosphoproteomics analysis by Liquid Chromatography Combined with Tandem Mass Spectrometry (LC-MS/MS)

Be(2)-C cells were cultured in triplicates in 10 cm dishes and treated the following day either with MitoQ EC50, DPI EC50, Vin EC50, (M+D)/2, (M+D+V)/3, or vehicle for 24 h and washed once with PBS. Cells were lysed in 100 μL ice cold 8 M Urea / 50 mM Tris HCl lysis buffer (Fisher Scientific) with 1:100 of Halt Cocktail Inhibitor, EDTA-free (100X) (Thermo Scientific). Samples were sonicated on ice twice for 9 seconds, at a power of 15% with a Skl-150IIDN Ultrasonic Homogenizer (Medical Supply Co. Ltd.), and protein concentration was measured using the Pierce BCA protein assay (Thermo Scientific). The same amount of protein (400 μg) were used for all samples, and 8 M Urea lysis buffer was used as a diluent if required. 8 mM of reducing agent DTT (Sigma) was then added to the samples and incubated in a thermomixer at 30 °C for 30 minutes, followed by the addition of 20 mM iodoacetamide (Sigma) for protein carboxylation with incubation at 30 °C in the dark for 30 minutes. Samples were diluted with 50 mM Tris-HCl to bring urea concentration down to below 2 M to prevent trypsin inhibition. Trypsin / Lys-C Mix, Mass Spec Grade (Promega) was added to the samples which were digested overnight at 37 °C, on a thermomixer. Protein digestion was terminated by adding 100% formic acid to 1% final volume. Peptides were extracted from each sample using HyperSep C18 tips (Thermo Scientific), and 50 μg and 350 μg were kept for total proteomics and phosphoproteomics analysis, respectively.

Phospho-peptide enrichment was performed using TiO_2_ (Titansphere Phos-TiO Bulk 10 um, GL Science), ratio of beads to sample 10:1, mixed for 30 minutes at room temperature in 80% acetonitrile, 6% trifluoroacetic acid, 5 mM monopotassium phosphate, 20 mg/mL 2.5-dihydroxybenzoic acid. The phosphopeptides were eluted by adding 80 μL of elution buffer directly (50% acetonitrile, 500 uL H_2_O, 7% Nh4OH ammonium hydroxide). Eluents were evaporated on the CentriVap Concentrator (Labconco) at 45 °C for an hour. Peptides and phosphopeptides were analyzed separately by mass spectrometry.

Samples were analyzed on a timsTof Pro Mass Spectrometer (Bruker) as described previously^84^.

The raw data was searched against the *Homo sapiens* subset of the Uniprot Swissprot database (reviewed) using the Fragpipe proteomics pipeline 20.0 with specific parameters for TIMS data-dependent acquisition (TIMS DDA). The normalized protein intensity of each identified peptide was used for label-free quantitation using the LFQ-MBR workflow. Within MSFraggers, precursor mass tolerance was set to +/-20ppm, fragment mass tolerance was set to +/-20ppm, tryptic cleavage specificity was applied, as well as fixed methionine oxidation and fixed N-terminal acetylation. Phospho-proteomic analysis was carried out using the LFQ-phospho workflow which includes phosphorylation as a variable modification on serine, threonine, and tyrosine residues^85^.

### 4.12. Proteomics and Phosphoproteomics Data Analysis Using Perseus

Analysis for treatment versus untreated control of peptides and phosphopeptides was performed using Perseus v2.0.9.0. Phosphoproteomics results were normalized to the total proteomics. The data file from the Fragpipe output was loaded as a generic matrix in the software, with the Max LFQ intensities as main columns. The rows were filtered to remove the reverse proteins, the proteins only identified by site and the contaminants. The values were log2 transformed into normally distributed data. Rows were filtered based on the number of valid values: a minimum of 4 valid values out of 6 replicates in at least one group. Then, missing values were imputed from normal distribution by calculating width and center of the distribution. Statistical analysis was done using a One-Sample t-Test (s0 (fold change) = 0.1, FDR = 0.05). Kinase Substrate Enrichment Analysis (KSEA) was performed using the fgsea R package. Kinase-substrate relationships were obtained from the PhosphoSitePlus database and provided as gene sets (<u>PhosphoSitePlus</u>). Further protein-protein interaction and pathway enrichment analysis were done by exporting the final matrix to STRING database.

### 4.13. Pathway Enrichment Analysis Using STRING

Upregulated and downregulated protein lists from each treatment condition versus untreated control were exported to the STRING database (https://string-db.org) for functional enrichment analysis and PPI network, using the k-means clustering method.

### 4.14. Linear Regression Model for Mechanism of Synergy of Drug Combinations

The Fragpipe combined_proteins.txt file was filtered to remove known contaminants and protein groups containing no protein. Protein LFQ intensity values were log base 2 transformed, with zero values being assigned NaN to circumvent undefined values. Filtering was applied to keep only those proteins with at least four valid values in at least one drug treatment condition. NaN values were then reassigned to zero before median centering. Site- and condition-specific imputation was performed using the scImpute function in PhosR^86^ if the protein was quantified in more than 50% of samples from the same drug treatment. Sample-wise tail-based imputation was implemented with the tImpute function^86^. Linear regression through limma was used to model the relationship between protein abundance and drug treatment. The design matrix was coded as model.matrix(∼treatment), with control samples representing the reference. Differential effects were identified by contrasting the coefficients from the combination drug treatment to the addition of each drug treatment in isolation. Differential protein abundance by empirical Bayes was performed to identify proteins reaching statistical significance. A signature of combination drug treatment synergy was created by selecting statically significant (adj p value < 0.05) and upregulated (LogFC > 0) genes from the above comparison. Gene set enrichment analysis (GSEA) was performed using the fgsea R package on the ontology gene sets retrieved from gsea-msigdb.org (https://www.gsea-msigdb.org/gsea/msigdb). An FDR threshold of 0.1 was implemented.

### 4.15. 3D Neuro-Spheres Imaging

Cells were seeded per well in 100 μL PrimNeuS media on a Nunclon Sphera 96-Well U-Shaped-Bottom Microplate (Thermo Scientific). Four days after seeding, the cells were treated with MitoQ EC50 + DPI EC50, MitoQ EC50 + DPI EC50 + Vin EC50, or vehicle for 24 h. The spheres were washed once with PBS, then fixed in acetone, washed twice with PBS, and permeabilized using Triton X-100. The spheres were washed twice and stained with 1:500 PI solution (Bio-Sciences) supplemented with 250 μg/mL RNAse A (Qiagen) and Hoechst nuclear counterstain (2 μg/mL) (Bio-Sciences) for 30 min protected from light, at room temperature. Spheres were then washed once in PBS and imaged in fresh PBS on an Olympus IX83 Inverted Microscope (Mason Technology) using the TRITC (Ex/Em: 396/580 nm) and DAPI (Ex/Em: 353/483) filter sets. Image processing was performed on ImageJ (v1.53e; Java 1.8.0_172).

### 4.16. Statistics

All normalized values were transferred to GraphPad Prism 8.4.3 for statistical analysis of biological replicates. All graphs are represented as mean ± SEM. Two-Way ANOVAs were used when using the IMR-5/75 and SH-SY5Y/MYCN -Dox/+Dox cell systems.

## Acknowledgements

This work received funding from Science Foundation Ireland and National Children’s Research Centre / Children’s Health Ireland through the Precision Oncology Ireland grant 18/SPP/3522, the Science Foundation Ireland Frontiers for the Future Program grant 22/FFP-A/10729, and with the financial support of Children’s Health Foundation and under the management of Science Foundation Ireland under the Frontiers for the Future Program Grant Number 21/FFP-P/10130. Mass spectrometry was performed at The Comprehensive Molecular Analytical Platform which was funded by The SFI Research Infrastructure Program (18/RI/5702). Real-Time PCR was performed at the Genomics Core facility of the UCD Conway Institute of Biomolecular and Biomedical Research. The Opera Phenix High Content Screening System (PerkinElmer) was used in the UCD Cell Screening laboratory, in UCD School of Biology and Environmental Science. LC and EP were funded by Cancer Research UK Discovery Programme Grant (A28278). LC was also funded by HEFCE/RAE.

## Author contributions statement

Conceptualization: S.E., W.K., and M.H.; original draft preparation: S.E.; review and editing: W.K. and M.H.; data analysis: S.E. and D.E.; mass spectrometry: K.W.; neurospheres generation and *in vivo* experiments: E.P. All authors have read and agreed to the published version of the manuscript.

