## Supplementary Materials for "Oxphos Targeting of *Mycn*-Amplified Neuroblastoma"

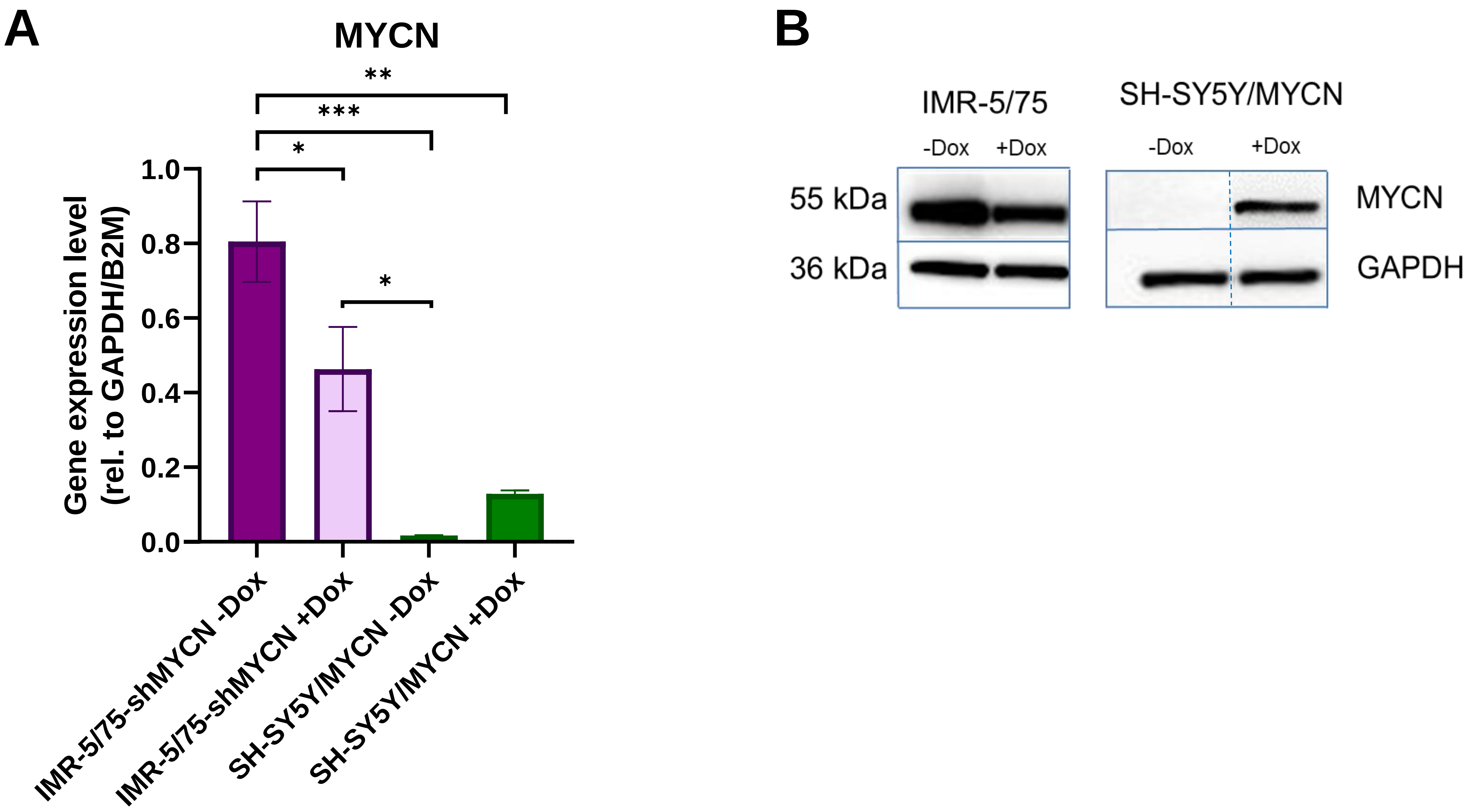


**Fig. S1. MYCN mRNA and protein expression level in the two MYCN-regulatable NB cell lines, IMR-5/75 (MNA) and SH-SY5Y/MYCN (non-MNA). A)** MYCN gene expression level was measured in IMR-5/75 and SH-SY5Y/MYCN by RT-qPCR after 24 hours plasmid induction with doxycycline treatment. Gene expression was normalized to two housekeeping genes, *GAPDH* and *B2M*. (Mean ± SEM; n=3; Ordinary One-Way ANOVA, $p<0.05$ = *, $p<0.005$ = **, $p<0.0005$ = ***, $p<0.0001$ = ****). **B)** MYCN protein expression level was measured in IMR-5/75 and SH-SY5Y/MYCN by Western-blotting after 24 hours plasmid induction with doxycycline treatment. The SH-SY5Y lanes were spliced from samples analyzed on the same blots as indicated by the broken line. Image is representative of 3 individual biological replicates.

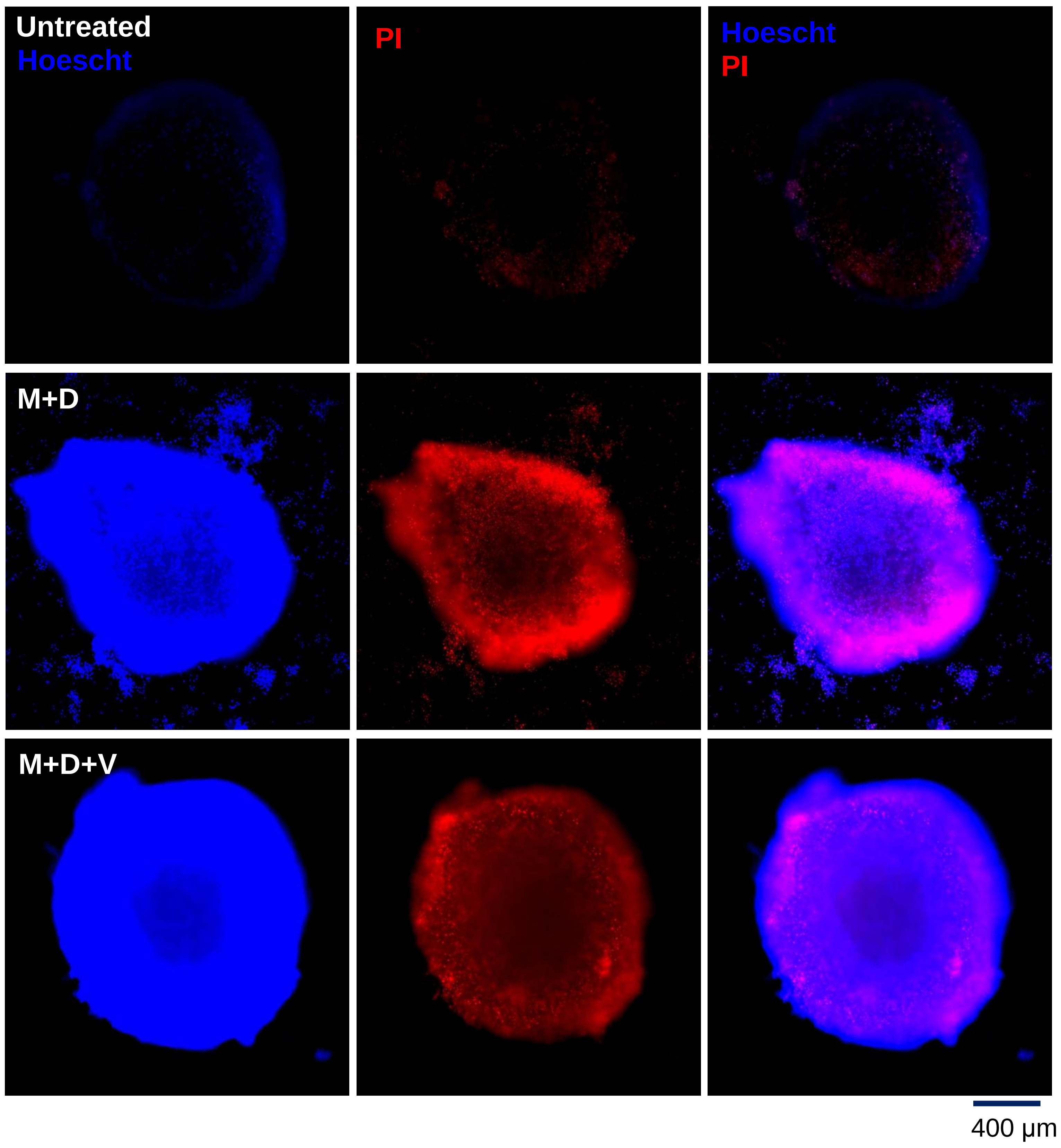


**Fig. S2. Double and triple combinations induce cell death in NB 3D sphere model.** Fluorescent microscopy images of Th-*MYCN* spheroids treated with control, MitoQ + DPI (EC50) or MitoQ + DPI + vincristine (EC50) for 24 hours (x10) and stained with Hoechst and Propidium Iodide. Scale bar: 400 μm. n=2.

**A**

| Description | Count in network | P-value enrichment | Proteins |
|---|---|---|---|
| **Histone H3-K27 and H3-K4 trimethylation** | 6 out of 50 | < 1.0e-16 | H1-0, H1-2, H1-3, H1-4, H1-5, H1-10 |
| **Carboxylic acid metabolic process** | 5 out of 50 | 3.7e-10 | ASL, ASNS, ASS1, GOT1, IDH1 |
| **Vesicle fusion** | 3 out of 50 | 6.88e-06 | STX10, VAMP7, VPS33A |
| **Hydrogen peroxide catabolic process** | 2 out of 50 | 3.7e-3 | HBA2, HBD |
| **Endoplasmic reticulum** | 4 out of 50 | 1.8e-3 | AHSG, FN1, SERPINH1, VTN |
| **Aminoacyl-tRNA ligase activity** | 2 out of 50 | 1.0e-2 | CARS1, WARS1 |

**B**

| Description | Count in network | P-value enrichment | Proteins |
|---|---|---|---|
| **Mitochondrial ATP synthesis coupled electron transport** | 17 out of 104 | < 1.0e-16 | NDUFA2, NDUFA5, NDUFA6, NDUFA7, NDUFA9, NDUFA12, NDUFB4, NDUFB9, NDUFS1, NDUFS2, NDUFS3, NDUFS6, NDUFS7, NDUFS8, NDUFV1, NDUFV2, UQCRC2 |
| **Mitochondrial ribosome/gene expression** | 16 out of 104 | < 1.0e-16 | MRPL13, MRPL14, MRPL21, MRPL22, MRPL3, MRPL34, MRPL37, MRPL47, MRPL9, MRPS14, MRPS21, MRPS30, MRPS33, MRPS35, MRPS5, PTCD3 |
| **Cell division, mitosis** | 15 out of 104 | < 1.0e-16 | AURKB, CCNB1, CKS1B, KIFF11, PLK1, PRC1, PTTG1, RACGAP1, RRM2, TK1, TYMS, TOP2A, UBE2C, UBA52, UBE2S, |
| **DNA replication** | 2 out of 104 | 2.2e-2 | ASF1B, CHAF1B |

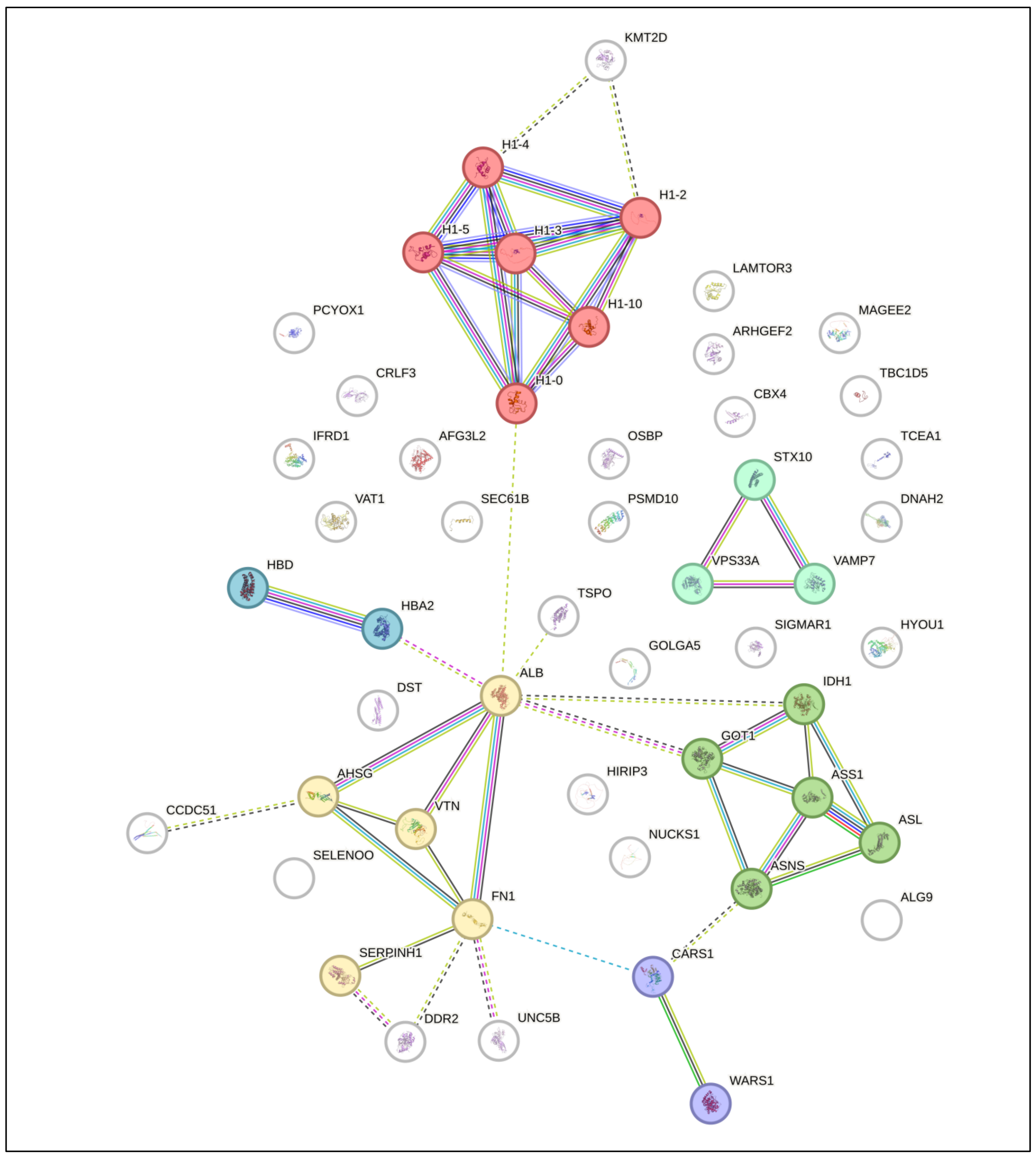


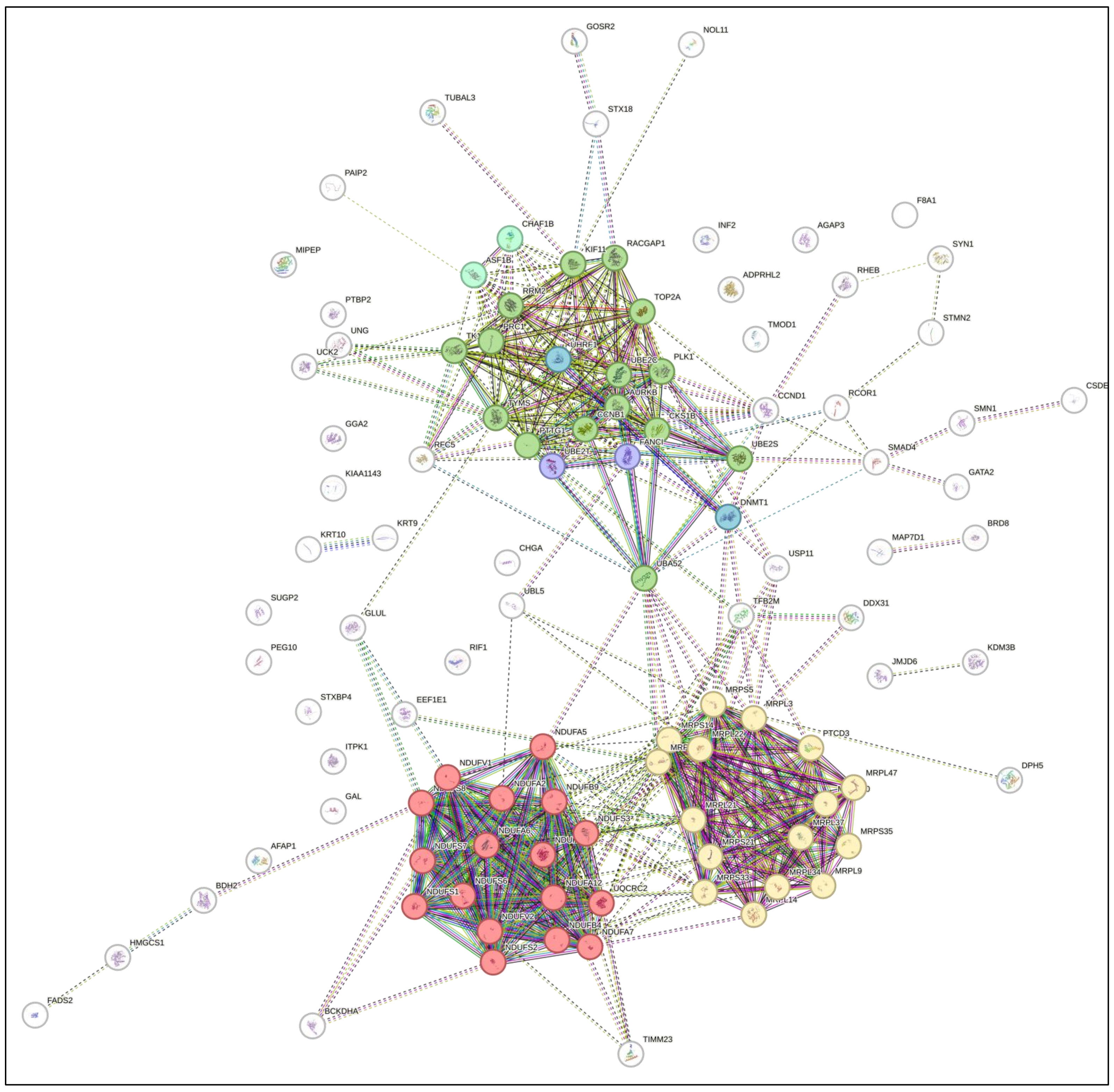


**Fig. S3. STRING visualization of significantly enriched proteins and associated pathways in Be(2)-C MNA cell line treated with MitoQ EC50.** Be(2)-C cells were treated with MitoQ EC50 for 24 hours and analyzed by mass spectrometry. **A)** Upregulated proteins and **B)** downregulated proteins compared to the control condition. (Mean ± SEM; n=3; Two-sample t-test, FDR = 0.05, s0 = 0.1)

**A**

| Description | Count in network | P-value enrichment | Proteins |
|---|---|---|---|
| **Cholesterol/small molecule metabolic process** | 3 out of 34 | 3.56e-05 | CYP51A1, HMGCR, SQLE |

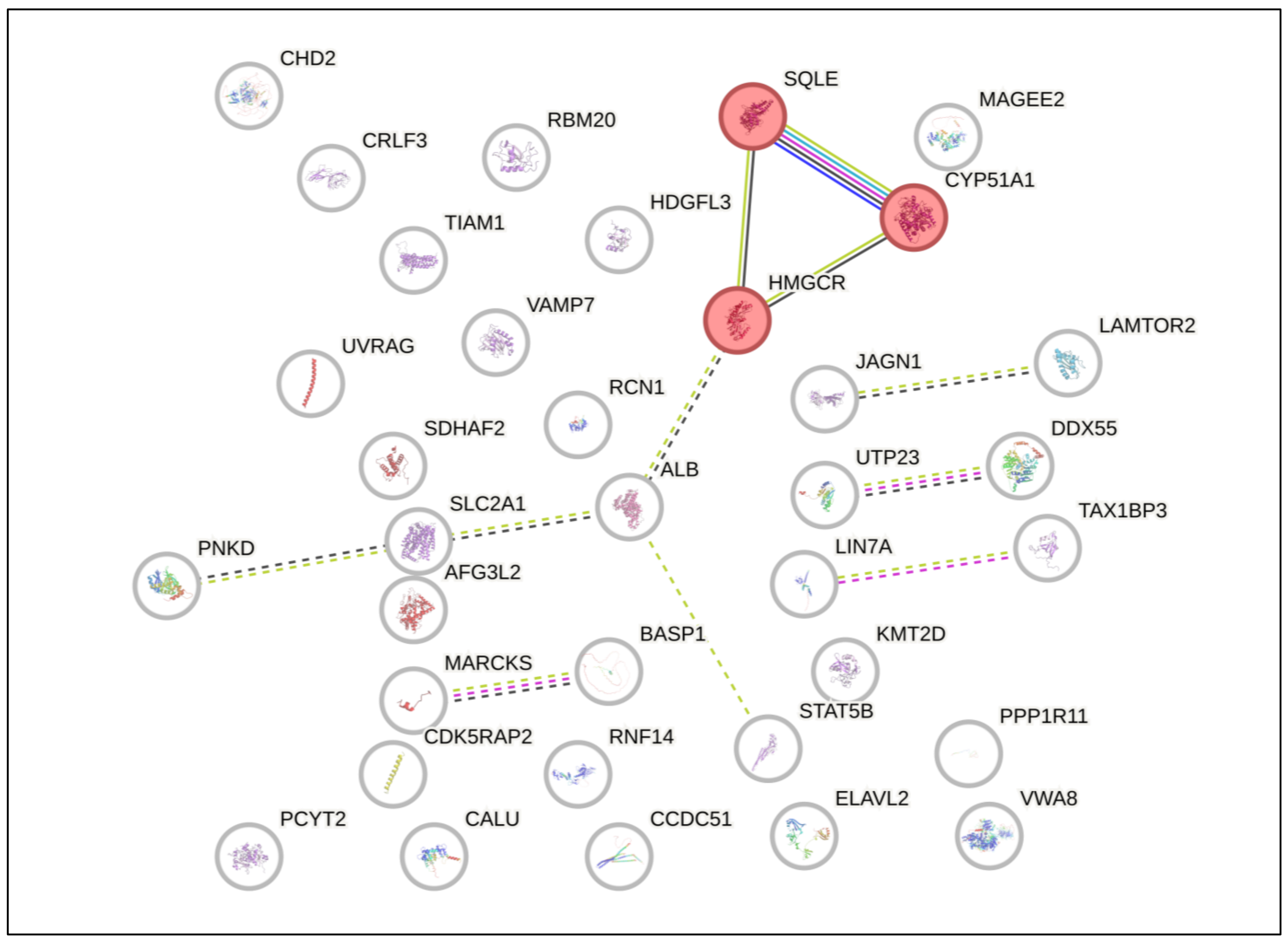


**B**

| Description | Count in network | P-value enrichment | Proteins |
|---|---|---|---|
| **Regulation of cell cycle** | 10 out of 33 | < 1.0e-16 | ASF1B, CCNB1, CENPF, CKAP2, KIF11, PCLAF, PRC1, RRM2, TOP2A, UBE2C |
| **Mitochondrial ATP synthesis coupled electron transport** | 10 out of 33 | < 1.0e-16 | MRPL34, NDUFA12, NDUFA13, NDUFA7, NDUFB6, NDUFS1, NDUFS6, NDUFS8, NDUFV1, TTC19 |
| **Histone H3-K27 and H3-K4 trimethylation, structural molecule** | 3 out of 33 | 5.71e-08 | H1-2, H1-3, H1-4 |
| **Iron-sulfur cluster assembly** | 2 out of 33 | 1.0e-3 | CIAPIN1, YAE1 |

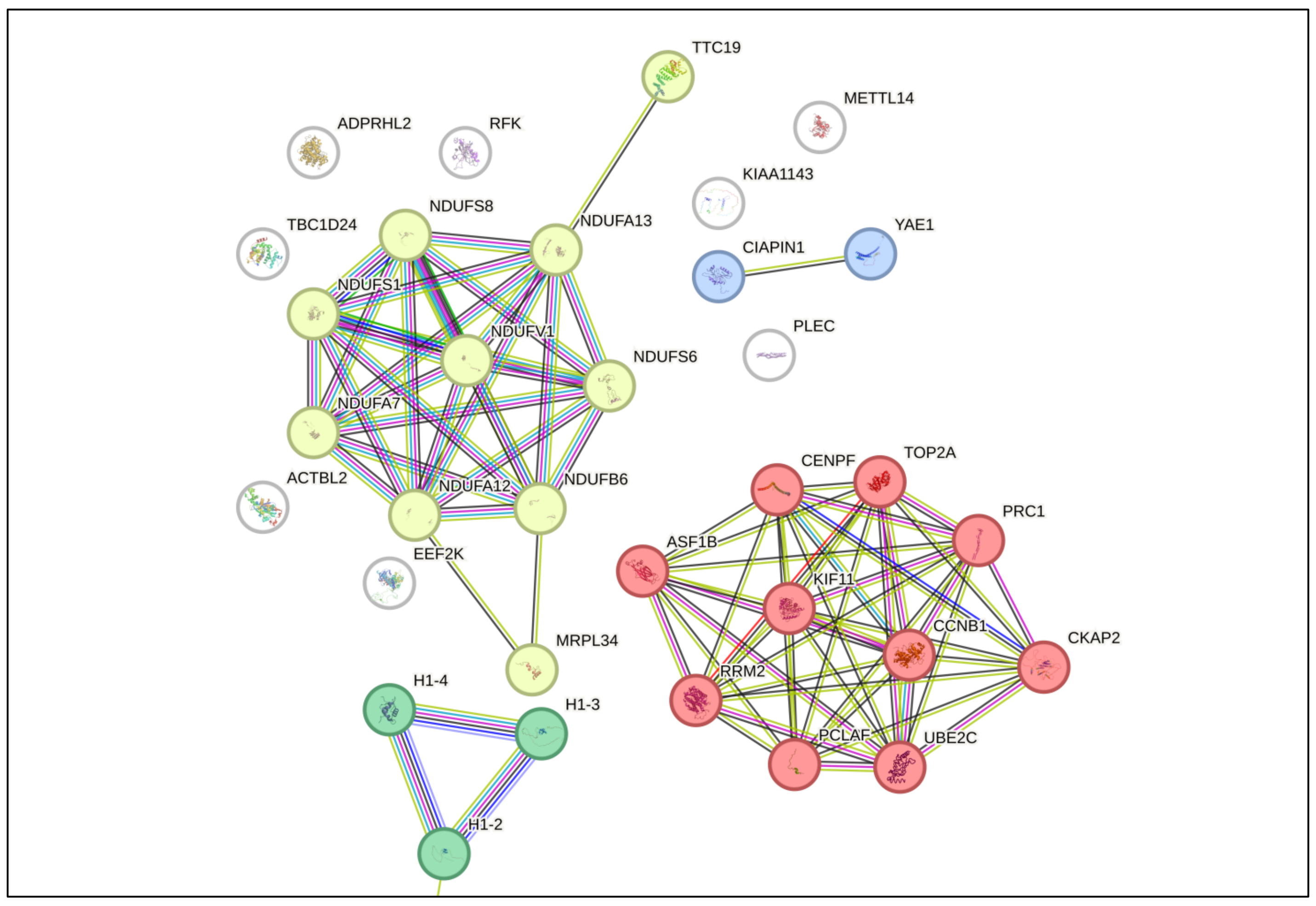


**Fig. S4. STRING visualization of significantly enriched proteins and associated pathways in Be(2)-C MNA cell line treated with DPI EC50.** Be(2)-C cells were treated with DPI EC50 for 24 hours and analyzed by mass spectrometry. **A)** Upregulated proteins and **B)** downregulated proteins compared to the control condition. (Mean ± SEM; n=3; Two-sample t-test, FDR = 0.05, s0 = 0.1)

A

| Description | Count in network | P-value enrichment | Proteins |
|---|---|---|---|
| **Cell division, mitosis** | 14 out of 27 | < 1.0e-16 | ASF1B, AURKB, CCNB1, KIF11, KPNA2, PCLAF, PLK1, RRM2, TK1, TOP2A, TUBA4A, TYMS, UBE2C, UBE2S |
| **Mitochondrial ATP synthesis coupled electron transport/ribosome** | 11 out of 27 | < 1.0e-16 | ATP5F1C, MFN2, MRPL21, NDUFA12, NDUFA6, NDUFA7, NDUFA9, NDUFS1, NDUFS6, NDUFS8, NDUFV1 |

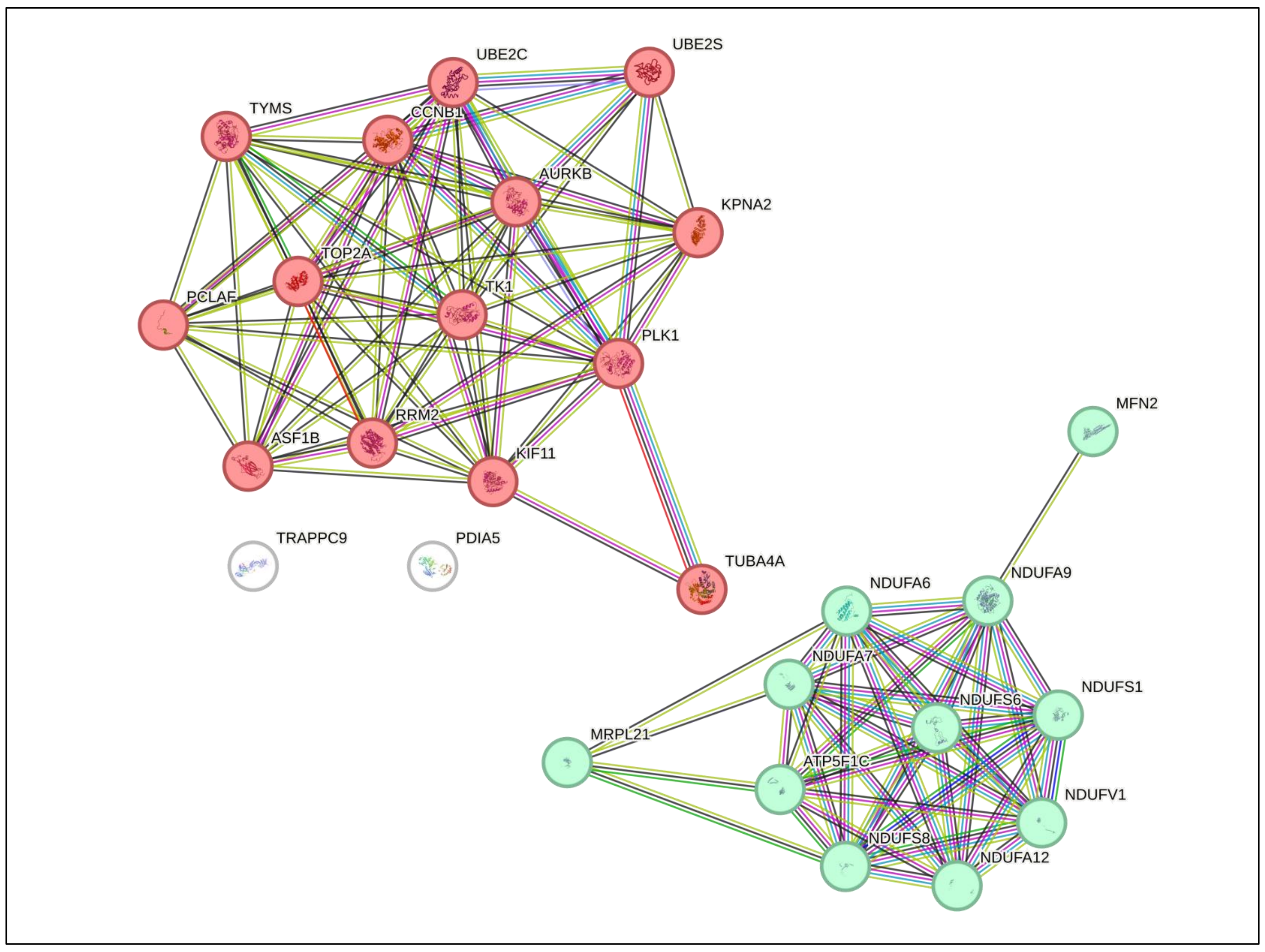


**Fig. S5. STRING visualization of significantly enriched proteins and associated pathways in Be(2)-C MNA cell line treated with (M+D)/2.** Be(2)-C cells were treated with (M+D)/2 for 24 hours and analyzed by mass spectrometry. **A)** Downregulated proteins compared to the control condition. (Mean ± SEM; n=3; Two-sample t-test, FDR = 0.05, s0 = 0.1)

**A**

| Description | Count in network | P-value enrichment | Proteins |
|---|---|---|---|
| **Mitochondrial ATP synthesis coupled electron transport** | 3 out of 5 | 9.77e-05 | NDUFA7, NDUFA12, NDUFS6 |
| **P53 signalling pathway/Cell cycle** | 2 out of 5 | 0.104 | CCNB1, RRM2 |

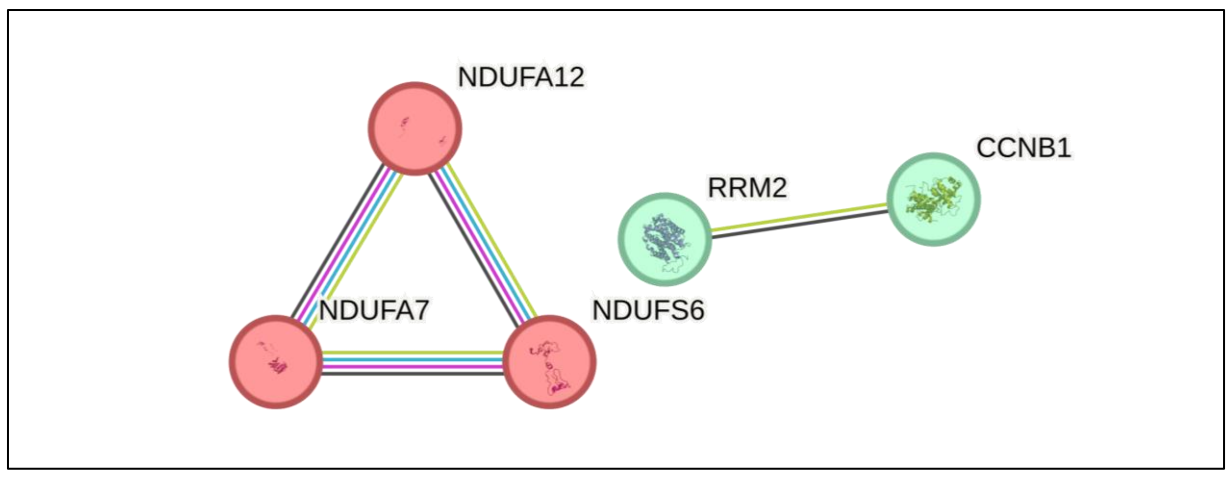


**Fig. S6. STRING visualization of significantly enriched proteins and associated pathways in Be(2)-C MNA cell line treated with (M+D+V)/3.** Be(2)-C cells were treated with (M+D+V)/3 for 24 hours and analyzed by mass spectrometry. **A)** Downregulated proteins compared to the control condition. (Mean ± SEM; n=3; Two-sample t-test, FDR = 0.05, s0 = 0.1)

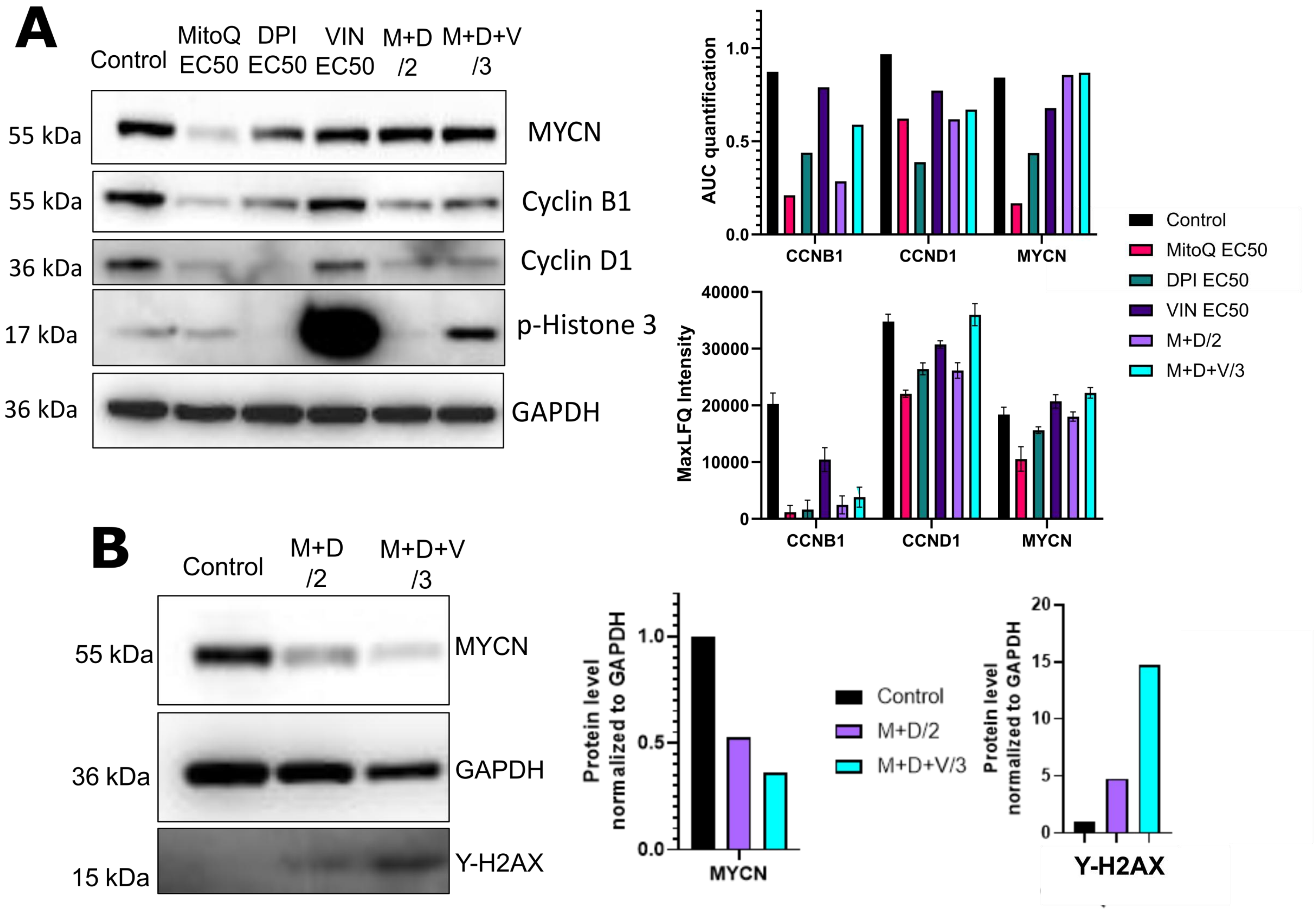


**Fig. S7. Effect of single and combination treatments on cell cycle protein markers and MYCN level. A)** Western blot analysis of cyclin B1 (CCNB1), cyclin D1 (CCND1), MYCN, and pS10-phospho histone 3 in Be(2)-C cells treated under the same conditions as in the proteomic analysis (n=1). The right upper panel shows the associated area under the curve (AUC). The right lower panel presents LFQ and normalized protein content values of CCNB1, CCND1, and MYCN as detected by MS-based proteomics (Mean ± SEM; n=3, two-sample t-test, $s_0$ = 0.1, FDR = 0.05).**(B)** Western blot analysis of MYCN and γ-H2AX in IMR-5/75 cells treated with M+D/2 or M+D+V/3 for 24 hours. GAPDH was used as a loading control (n=1).

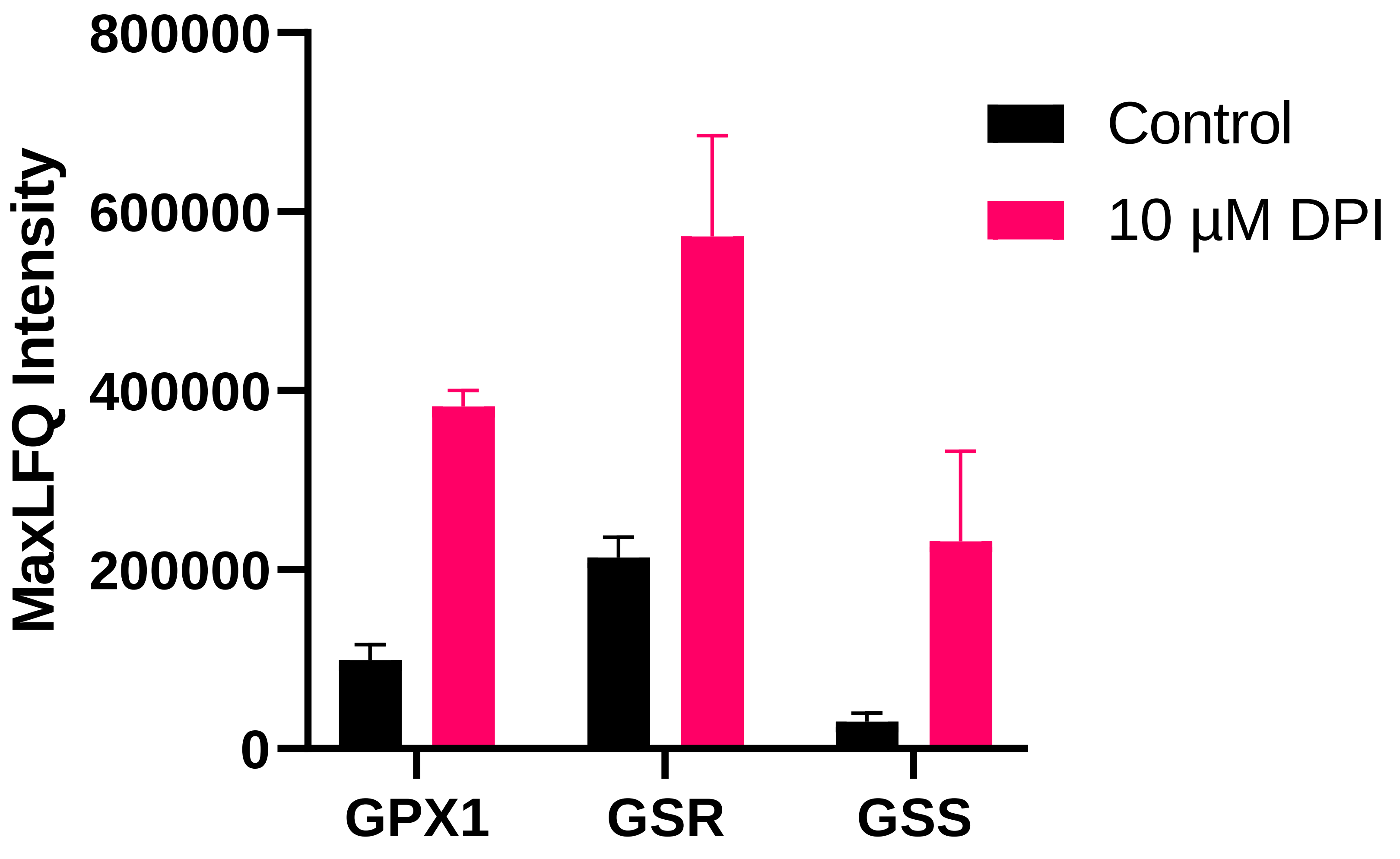


**Fig. S8. DPI increases key antioxidant protein levels in Be(2)-C cells.** Max LFQ values of Glutathione Peroxidase 1 (GPX1), Glutathione Reductase (GSR) and Glutathione Synthetase (GSS) in Be(2)-C as detected by MS based proteomics under 10 μM DPI treatment. (Mean ± SEM; n=3, Two-sample t-test, s0 = 1, FDR = 0.01).
